# Rapid enrichment of progenitor exhausted neoantigen-specific CD8 T cells from peripheral blood

**DOI:** 10.1101/2025.05.11.653315

**Authors:** Mamduh Khateb, Raina Jung, Stav Leibou, Patrick Hadley, Zhiya Yu, Aaron J. Dinerman, Victoria Dulemba, Billel Gasmi, Noam Levin, Peter Kim, Aarushi Bhasin, Deepali Bhat, Sivasish Sindiri, Jared J. Gartner, Todd D. Prickett, Tiffany Benzine, Shahram S. Farid, Maria R. Parkhurst, Hyunmi Halas, Yaqiang Cao, Keji Zhao, James C. Yang, Paul F. Robbins, Frank Lowery, Sri Krishna, Theo Heller, Daniel McVicar, Steven A. Rosenberg, Nicholas D. Klemen

**Affiliations:** Surgery Branch, NCI, NIH, Bethesda, MD USA; Systems Biology Center, NHLBI, NIH, Bethesda, MD USA; Liver Diseases Branch, NIDDK, NIH, Bethesda MD USA; Cancer Innovation Laboratory, NCI/CCR, NIH, Frederick MD USA

## Abstract

Neoantigen-reactive peripheral blood lymphocytes (NeoPBL) are tumor-specific T cells found at ultra-low frequencies in the blood. Unlike tumor-infiltrating lymphocytes (TIL), NeoPBL exist in a favorable less dysfunctional phenotypic state *in vivo*, but their rarity has precluded their effective use as cell therapy. Leveraging *a priori* knowledge of bona fide neoantigens, we combined high-intensity neoantigen stimulation with bead extraction of neoantigen peptide-pulsed target cells to enable the enrichment of NeoPBL to frequencies comparable to *ex vivo* cultured TIL over a 28-day period. Throughout this process, NeoPBL demonstrate specific reactivity against autologous tumor organoids and maintain memory-like features, including elevated expression of *CD28* and *TCF7*. We additionally demonstrate that NeoPBL reactivity is polyclonal, encompassing multiple clonotypes that are detectable within *in vivo* TIL populations, underscoring physiological specificity for the targeted neoantigens. This streamlined process yields clinically relevant cell doses and enables identification and expansion of blood-derived neoantigen-specific TCRs. By potentially avoiding additional surgical risks and protracted delays of TIL and individualized TCR-engineered methods, the NeoPBL platform may have clinical and practical advantages. Ultimately, NeoPBL combines intrinsic cell fitness, minimal invasiveness and rapidity to potentially facilitate personalized adoptive cell therapy for cancer.

## Introduction

Cancer remains one of the most pressing global health challenges, claiming millions of lives globally each year and demanding new, more effective immunotherapies. Within this urgent landscape, metastatic gastrointestinal (GI) malignancies – especially pancreatic ductal adenocarcinoma (PDAC) and colorectal cancer (CRC) – stand out for their lethality, causing over 100,000 deaths per year in the United States alone^1,2^. We recently reported that tumor-infiltrating lymphocytes (TIL) or TCR-engineered peripheral blood lymphocytes (TCR-T cells) can mediate regression of metastatic mismatch repair–proficient GI cancers, demonstrating the therapeutic potential of adoptive T cell transfer^3,4^.

Despite the success of TIL in melanoma, where complete responses (CRs) are frequent and durable^5,6^, its application in GI cancers has mostly yielded partial regressions^4^. One explanation for these results is that TIL isolated from GI tumors exist in a highly differentiated, dysfunctional state^7,8^. TCR-T cell therapy was devised to circumvent this problem by cloning neoantigen-specific TCRs (neoTCRs) into peripheral blood lymphocytes (PBL), which are typically far less differentiated than TIL^9^. Judging by the high proportion of transferred TCR-T cells with a stem-like CD39^-^CD69^-^ phenotype^3,7^, this strategy may be a step towards resolving the problem of advanced TIL differentiation, but so far TCR-T cells have only rarely mediated CRs.

These outcomes point to the limitations of using a monoclonal attack on tumors, which are antigenically heterogeneous. A key limitation of TCR-T cell therapy is its focus is often a single antigen, enabling tumor escape via antigen or HLA loss^3^. From an immune perspective, polyclonal T cell responses against each targeted neoantigen could also potentially improve anti-tumor responses. A clinical demonstration of this comes from a patient with CRC who had long-term disease control after infusion of a TIL product recognizing a KRAS G12D neoantigen^10^. Follow-up analyses uncovered several distinct KRAS G12D-specific T cell clonotypes, each displaying a unique affinity and functional avidity profile^11^. The frequency at which conventional TIL cultures generate similarly polyclonal repertoires, however, remains unclear.

Studies of patients with melanoma treated with immune checkpoint blockade also showed deep clinical responses occurred when there were multiple T cell clonotypes recognizing a few immunodominant mutations^12^. These results suggest that T cells with different avidities could have distinct functional roles in the tumor microenvironment. Engaging both high- and intermediate-avidity T cells might produce more sustained anti-tumor effects because TCR signal strength is a key determinant of T cell responses and tumor escape^13^. Collectively, these data highlight that there could be advantages of mounting polyclonal T cell responses against each bona fide neoantigen.

TIL and TCR-T cell therapy also have important clinical and practical limitations. TIL usually requires an extra surgical procedure and typically cannot commence until metastases appear. Personalized TCR-T cell therapy is expensive, technically demanding and can take months of development for targeting bespoke patient unique antigens – too long for many patients with advanced GI cancer. “Off-the-shelf” TCR-T cell therapy precisely matched for HLA and directed against a shared tumor antigen is much faster, yet HLA genes are among the most diverse in the human genome, so many of these treatments will necessarily be monoclonal^14–17^.

An alternative source of autologous lymphocytes for cell therapy is the peripheral blood. *In vitro* sensitization (IVS) has previously been used to raise PBL specific for melanoma antigens^18–20^. More recently, attention has turned to autologous patient-derived neoantigen-specific peripheral blood lymphocytes (NeoPBL) as a platform for ACT, as they potentially combine minimal differentiation, polyclonality and neoantigen specificity^21^. The challenge with NeoPBL is their ultra-low frequency, estimated to be fewer than 5 per 100,000 cells, which has impeded their reliable detection, expansion and enrichment to clinically relevant levels^22^. Consequently, attempts to utilize NeoPBL for cell therapy has resulted in a very low frequency of NeoPBL in the infused product, perhaps accounting for the limited evidence of clinical efficacy so far^21^.

To rapidly generate large numbers of T cells in a predictable time scale, adoptive T cell therapies commonly rely on a rapid expansion protocol (REP) which uses anti-CD3 (OKT3) stimulation plus irradiated feeder cells in cytokine-rich media. Important downsides of a REP are that it causes T cell differentiation and can result in overgrowth of bystander cells because OKT3 induces nonspecific T cell activation. However, recent work describing *NeoExpand* showed *in vitro* sensitization (IVS) using neoantigen peptides can enrich for rare TIL reactivities while improving the phenotype of expanded cells, opening the door to new approaches to isolate and expand neoantigen-specific T cells^23^.

Guided by the goal of generating a highly enriched, minimally differentiated and rapidly available T cell product, we developed NeoSelect: a protocol that merges on target neoantigen stimulation, bead extraction of neoantigen peptide-pulsed target cells and OKT3-mediated expansion as in a REP. This culture-based approach results in the enrichment of NeoPBL to frequencies comparable to *ex vivo* cultured TIL over a 28-day period, while achieving large scale expansion and preserving key memory features and clonal diversity – without requiring an additional surgical procedure if targeted neoantigens are known.

## Results

### Neoantigen peptide stimulation prior to a Rapid Expansion Protocol (REP) increases blood-derived TCR-T cell expansion and enriches for a less differentiated T cell phenotype

Because NeoPBL are low in both frequency and absolute numbers, their clinical application may require an approach that both maximizes T cell expansion and reactivity enrichment for use in ACT^21^. To this end, we merged on target stimulation with a REP^23^. This unconventional *NeoExpand-REP* protocol begins with antigen-specific stimulation achieved through co-culture with irradiated autologous lymphocytes pulsed with predicted minimal class I neoantigen peptides for 16 hours, resulting in a high degree of T cell activation as indicated by 4-1BB expression **(Fig. 1A, S1A)**. After this, cells are transferred to REP conditions (with OKT3 and irradiated feeders) and expanded over the next 14 days.

**Figure 1.**
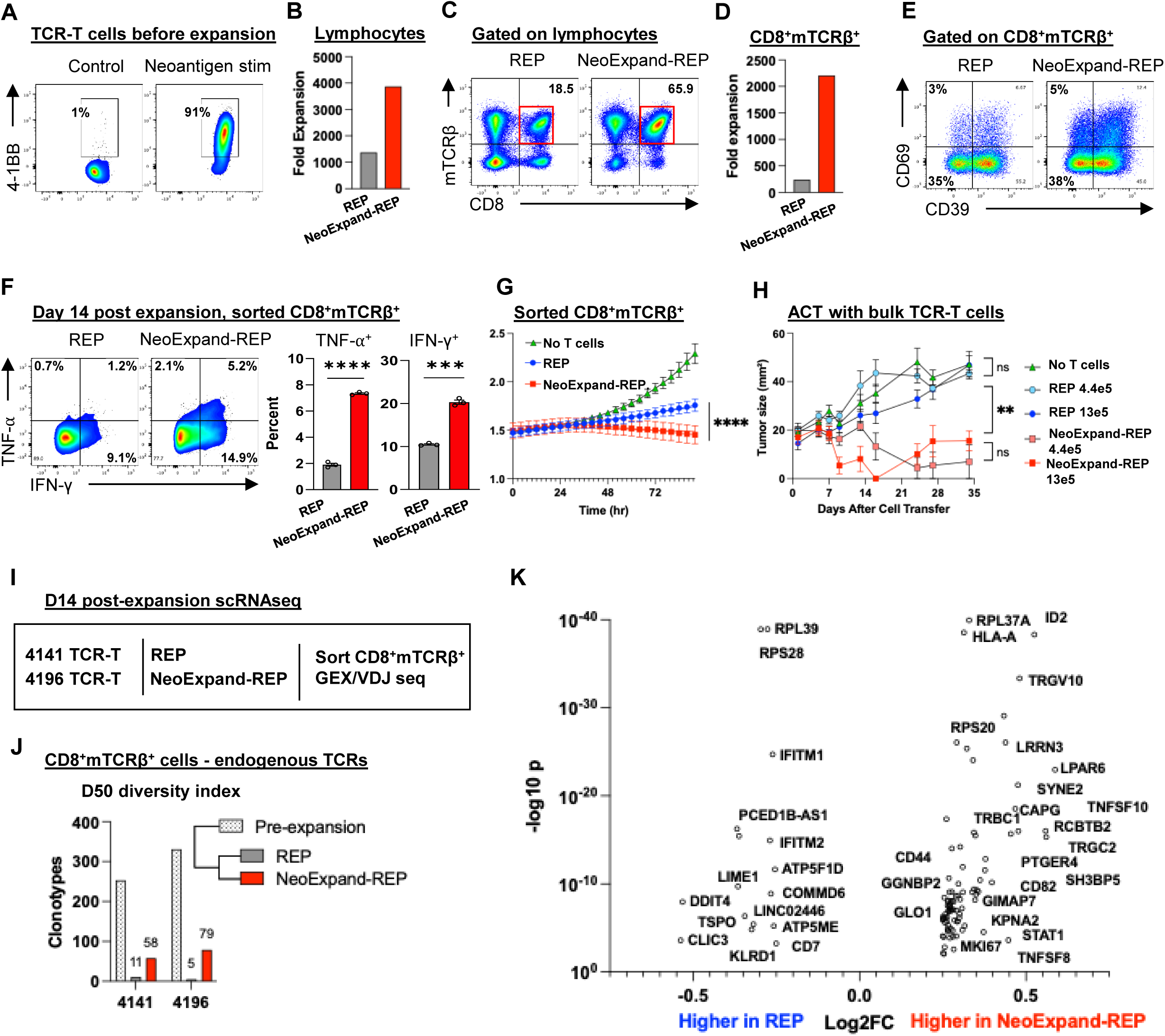
NeoExpand-REP enhances expansion, preserves diversity, and improves phenotype and function of TCR-T cells. (A) TCR-T cells targeting TP53 R175H (HLA-A*02:01-restricted) were generated from patient PBL were plated alone (control) or at a 1:5 E:T ratio with neoantigen peptide-pulsed PBL (neoantigen stim). Flow cytometry plots show 4-1BB expression on CD8^+^mTCRβ^+^ gated cells after 16 hours, after which cells were transferred to REP conditions (with OKT3, allogeneic feeders and IL2). (B) Fold expansion of all lymphocytes (CD4⁺, CD8⁺, transduced or untransduced) from 4196 TCR-T cultures after 14 days in rapid expansion protocol (REP) or NeoExpand-REP. (C) Flow cytometry data showing CD8 and mTCRβ expression on lymphocytes and (D) fold expansion of CD8^+^mTCRβ^+^ T cells during REP or NeoExpand-REP. **(E)** Phenotypic characterization of CD8⁺mTCRβ⁺ T cells by flow cytometry showing expression of CD39 and CD69. (F) Functional cytokine responses (TNF-α and IFN-γ) in sorted CD8⁺mTCRβ⁺ T cells following 16-hour co-culture with HLA-A*02:01^+^ TYK-nu tumor cells; bar graphs show technical replicates. (G) Tumor cell killing by sorted CD8⁺mTCRβ⁺ T cells (equal cell numbers) measured by luminescence in an Incucyte assay. (H) Tumor growth in NSG mice bearing established TYK-nu xenografts treated with either 4.4×10⁵ or 1.3×10⁶ TCR-T cells expanded via REP or NeoExpand-REP. Mice received one intravenous and two intraperitoneal doses of IL-2. Tumors were measured weekly by a blinded investigator. (I) Schematic of scRNA-seq workflow: TCR-T cells (from patients 4141 and 4196, both targeting TP53 R175H, with low and high avidity respectively) were expanded for 14 days via REP or NeoExpand-REP, sorted for CD8⁺mTCRβ⁺ expression, and captured for sequencing. (J) TCR diversity shown by D50 index—the number of clonotypes contributing 50% of the repertoire. Values for 4141 and 4196 TCR-T cells before and after expansion are shown. (K) Volcano plot showing differentially expressed genes (DEGs) in the top 5 expanded clonotypes present in both REP and NeoExpand-REP using a combined data set with both 4141 and 4196 TCR-T cells.

We first tested NeoExpand-REP using TCR-T cells as a model system with a known neoantigen target. We co-cultured equal numbers of TCR-T cells with autologous apheresis that had been pulsed with cognate minimal neoantigen peptide at effector to target (E:T) ratios of 5:1, 1:1 and 1:5 for 16 hours, after which they were transferred to a standard REP. NeoExpand-REP enriched TCR-T cells in a target cell dose dependent manner relative to a standard REP while also achieving large scale expansion **(Figs. S1A, S1B)**. Surprisingly, despite increasing expansion of TCR-T cells, NeoExpand-REP maintained or increased the fraction of TCR-T cells that were with a stem-like CD39^-^CD69^-^ phenotype, suggestive of a less differentiated phenotype^7^ **(Fig. S1B, S1C)**. These counter-intuitive results suggest NeoExpand-REP may be selecting and enriching for memory T cells, which are predicted by models to have greater therapeutic potential^24,25^.

### NeoExpand-REP enhances the functional potency of resulting TCR-T cells *in vitro* and *in vivo*

To understand the functional consequences of NeoExpand-REP, we evaluated TCR-T cells targeting a TP53 R175H neoantigen that were transduced under good manufacturing practices (GMP) and expanded at clinical scale **(Fig. 1A**, **1B)**. These cells expressed TCRs with murine constant chains that can be detected using a murine TCRβ antibody^26^. NeoExpand-REP increased the proportion of CD8^⁺^mTCRβ^⁺^ cells nearly 10-fold **(Fig. 1C**, **1D)** while maintaining the CD39^-^CD69^-^ fraction **(Fig. 1E)**. On a per-cell basis, cells expanded this way demonstrated enhanced cytokine responses **(Fig. 1F)** and *in vitro* tumor killing **(Fig. 1G)**. Finally, adoptive transfer of bulk (unsorted) NeoExpand-REP cells *in vivo* resulted in regression of established TYK-nu (human ovarian cancer) tumors expressing the cognate TP53 R175H epitope and HLA-A*02:01^27^ **(Fig. 1H)**. Similar effects were seen in TCR-T cells expressing a lower avidity TP53 R175H-specific TCR^27,28^ **(Fig. S1D-S1G)**. Together these results indicate that NeoExpand-REP increases the absolute number of TCR-T cells while preserving or enhancing their phenotypic and functional quality.

### NeoExpand-REP enhances TCR-T cell clonal diversity

To understand at the clonal level why NeoExpand-REP maintained anti-tumor potential during expansion relative to a REP, we assessed the clonotypic diversity by performing single-cell RNA sequencing (scRNA-seq) on expanded CD8^⁺^mTCRβ^⁺^ cells and tracked clonotypes using endogenous TCRs. This revealed that NeoExpand-REP preserved more clonal diversity compared to a REP, although both lost diversity relative to the starting pool **(Fig. 1I, J)**. This result suggests that the proliferative and phenotypic advantages of NeoExpand-REP could potentially be accounted for by maintenance of additional younger TCR-T cell clones.

To evaluate whether NeoExpand-REP altered the intrinsic transcriptional program of resulting TCR-T cells, we compared the gene expression profiles of top expanded clonotypes present in both conditions after 14 days of expansion **(Figure 1K)**. NeoExpand-REP cells exhibited elevated expression of several notable genes, including ID2, which antagonizes E-protein activity and helps sustain effector and memory CD8^⁺^ T-cell lineages^29^. GIMAP7 – linked to improved survival and greater immune infiltration – was also elevated^30,31^. Increased TNF, TNFSF8 (CD30L), and TNFSF10 (TRAIL) transcripts signaled an activated phenotype with preserved cytotoxic and immunomodulatory potential. Elevated CD82 and SYNE2, both involved in cytoskeletal remodeling and immunological-synapse formation^32^, may point to superior cellular fitness.

Genes expressed at lower levels in NeoExpand-REP cells including *PCED1B-AS1*, *TSPO*, and *DDIT4* do not appear to belong to a single unified pathway or transcriptional program directly linked to T cell differentiation, but are associated with stress responses and metabolic regulation, which can impact the functional state of T cells. In summary, these modest changes suggest a transcriptional program of early effector differentiation while preserving memory potential and survival, although the precise mechanism for this requires further elucidation.

### NeoExpand-REP enables expansion and functional preservation of neoantigen-specific TIL in an advanced state of differentiation

TCR-T cells are manufactured from PBL by insertion of TCR genes, while neoantigen-specific TIL are usually highly differentiated and therefore may be less effective on a per-cell basis at mediating tumor regression^4,8,33^. Previously, NeoExpand was shown to selectively expand rare TIL clones without detrimentally impacting their phenotype^23^. We therefore sought to assess if NeoExpand-REP could amplify anti-tumor TIL. We anticipated that TIL would be especially susceptible to activation-induced cell death (AICD) during high-intensity protocols such as NeoExpand-REP, since they would have fewer stem-like clones available to expand. We tested this using TIL from patient 4610, grown from a colorectal cancer lung metastasis. Cultures from several independent TIL fragments contained clonally expanded CD8^⁺^T cells expressing TCRvβ17, with reactivity specific for a neoantigen in *FBXO33* **(Fig. S2A–C)**. Their near-universal CD39 expression marked them as being highly differentiated TIL **(Fig. S2D)**.

NeoExpand-REP likewise boosted both the absolute number and proportion of CD8^⁺^vβ17^⁺^ T cells relative to the conventional REP **(Fig. S2E–H)**. The enrichment was striking when the starting vβ17^⁺^ frequency was ≤10 % **(Fig. S2G)**. Importantly, expression of CD39, PD-1, and LAG-3 were stable following expansion **(Fig. S2I, S2J)**, again showing NeoExpand-REP did not further exacerbate dysfunctional phenotypes. To assess function, we challenged expanded TIL with neoantigen peptide stimulation and found fully intact TNF-α and IFN-γ responses **(Fig. S2K)**.

To understand how NeoExpand-REP might impact the transcriptional state of TIL, we analyzed gene expression of the CD8^+^vβ17^+^ clonotype after performing scRNAseq on lymphocytes expanded in both conditions **(Fig. S2L).** NeoExpand-REP caused a relative shift away from a chronic activation–induced state and toward one of functional fitness, costimulation, and renewal. This was evident in the coordinated downregulation of a suite of interferon-stimulated genes (ISGs) and stress-related transcripts – *IFI44L*, *IFI27*, *MX1*, *MX2*, *ISG15*, *OAS1*, *USP18*, *RSAD2*, and *PLSCR1*. These genes are hallmark signatures of persistent type I interferon signaling, a pathway often associated with T cell exhaustion and chronic inflammation.

By contrast, NeoExpand-REP resulted in the modest upregulation of several genes associated with memory, costimulation and effector coordination. Chief among these is *CD27*, a TNF receptor superfamily member that can supports T cell survival, expansion and effector function that can be expressed in TIL after *CD28* expression is lost^24,34–36^ **(Fig. S2L)**. *KLRK1*, encoding the activating receptor NKG2D, further contributes to a cytolytic, tissue-patrolling phenotype in CD8^⁺^TIL that lack CD28 but retain robust effector function^37^. *TIMP1*, secreted by activated T cells, can enhance MHC-I expression on antigen-presenting cells^38^. *GIMAP7*, discussed earlier in the TCR-T cell context, was also upregulated.

In addition to TIL from patient 4610, we also tested NeoExpand-REP using neoantigen-specific TIL from different patients and found the expansion benefit for reactive cells was highly reproducible **(Fig. S2M)**. Taken together, the initial on-target stimulation of NeoExpand-REP **(Fig. S2E)**, followed by OKT3-supported expansion, may result in the maintenance of the clonal repertoire and the phenotypic state of TIL, while simultaneously increasing the frequency and absolute number of neoantigen-reactive cells in TCR-T and TIL settings.

### NeoSelect as a tool to enable rapid antigen-specific expansion of low frequency T cells

There has long been interest in utilizing blood-resident T cells for cell therapy due to the clinical advantages of non-invasive sample procurement and ability to obtain large amounts of starting material via pheresis^21^. Having established that NeoExpand-REP could efficiently expand and enrich neoantigen-specific TIL, we turned our attention to the challenge of amplifying NeoPBL from peripheral blood. To overcome the ultra-low frequency of circulating NeoPBL, we incorporated an antigen-specific T cell extraction technique at the end of NeoExpand-REP that confers a 5-20-fold enrichment multiplier **(Fig. 2A)**. We henceforth refer to this protocol as NeoSelect.

**Figure 2.**
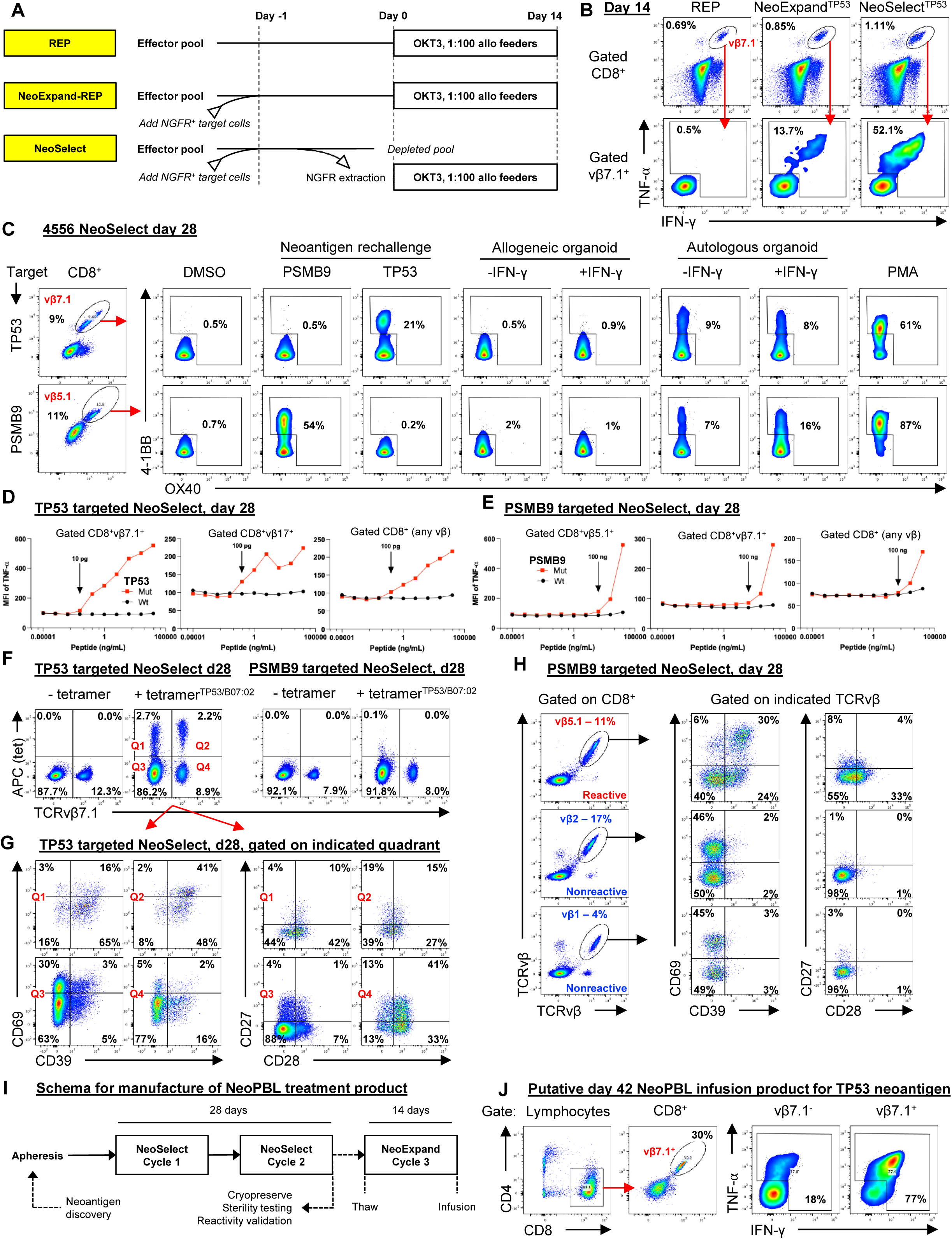
NeoSelect rapidly expands rare NeoPBL to clinically relevant frequencies. (A) Schematic comparing a standard REP, NeoExpand-REP and NeoSelect. (B) Representative flow cytometry showing the vβ7.1^+^ fraction of CD8^+^ gated T cells (top panels, gated CD8^+^) and cytokine responses (bottom panels, gated on CD8⁺vβ7.1⁺ T cells) of TP53-targeted NeoSelect cultures on day 14 to peptide rechallenge. (C) 4-1BB and OX40 expression on day 28 CD8⁺ T cells following 16-hour rechallenge with neoantigen peptides or tumor organoids. Gated on vβ7.1⁺ (TP53) or vβ5.1⁺ (PSMB9). **(D–E)** Peptide titration curves using mutant and wild-type peptides for TP53 (D) and PSMB9 (E); gating as indicated. (F) TP53/B*07:02 tetramer staining and vβ7.1 expression in day 28 cultures, gated on CD8⁺ T cells. (G) CD39 and CD69 expression (left) or CD27 and CD28 expression (right) in tetramer^+/-^ β7.1^+/-^ quadrants from TP53-targeted day 28 NeoSelect cultures. (H) CD39/CD69 and CD27/CD28 expression in CD8⁺ T cells gated on vβ5.1⁺ (predominantly PSMB9-reactive) versus non-reactive vβ families. (I) Schema for manufacturing a NeoPBL treatment product. This process is modeled based on TIL workflows, which require 25-30 days of *ex vivo* culture and are later expanded for 14 days to create an infusion product. (J) Expression of CD8 and vβ7.1, and cytokine responses (TNF-α, IFN-γ) following neoantigen rechallenge, of day 42 NeoPBL infusion product.

NeoSelect utilizes autologous CD4 T cells expressing truncated nerve growth factor receptor (NGFR) and pulsed with minimal neoantigen short peptides as target antigen presenting cells (APCs). After 16 hours of antigen-specific stimulation through co-culture with minimal peptide-pulsed target cells, an NGFR^+^ bead selection is used to extract target cells – which carry conjugated reactive effector cells along with them. The extracted NGFR^+^ pool is then expanded in REP conditions for 14 days. One cycle of NeoSelect is capable of exceeding 100-fold enrichment **(Fig. S3A, top panel)**, although the efficiency drops as reactive cell frequency approaches 1 **(Fig. S3A bottom panel)**. When two NeoSelect cycles are performed in series, they are multiplicative – potentially resulting in over 2,000-fold enrichment in 28 days, enough in theory to amplify NeoPBL from a frequency of 0.005% up to 10%.

We first evaluated NeoSelect in patient 4556, an individual with metastatic colorectal cancer. *Ex vivo* TIL cultures had failed to yield CD8^+^ neoantigen specific TIL, but neoTCRs predicted by TIL gene expression signatures revealed reactivity to TP53 I254T and PSMB9 R111Q neoantigens^8^. The neoTCRs expressed several different *TRBV* genes including TRBV4-1 (TCRvβ7.1) and TRBV5-1 (TCRvβ5.1). We then performed an experiment using diluted neoTCRs in a pool of untransduced cells that confirmed NeoSelect could mediate enrichment via the TP53 and PSMB9 minimal neoantigen peptides pulsed onto autologous target cells **(Fig. S3B)**.

Next, we obtained 3×10^8^ peripheral blood mononuclear cells from patient 4556 and performed CD8 memory bead selection to remove naïve CD8 T cells, reducing the risk of raising *de novo* CD8 reactivities. The baseline frequencies of CD8^+^ T cell clonotypes expressing TCRvβ7.1 and TCRvβ5.1 were each less than 2% **(S3C)**. We generated independent NeoSelect cultures, using the TP53 and PSMB9 neoantigen peptides separately. After the first cycle of NeoSelect (on day 14), TP53 reactivity was detectable in a growing CD8^+^vβ7.1^+^ compartment **(Fig. 2B, S3D)**. We did not detect TP53 reactivity outside of CD8^+^vβ7.1^+^ on day 14 **(Fig. S3E)** nor did we detect PSMB9 reactivity at that time point **(Fig. S3D)**.

At the beginning of the second NeoSelect cycle, immediately following bead extraction, we measured a 10-fold increase in activated (4-1BB^+^) vβ7.1^+^ T cells compared to no bead extraction, suggesting antigen-specific T cell extraction **(Fig. S3F)**. After two additional weeks of expansion, on day 28, the TCRvβ7.1^⁺^fraction of TP53-targeted cultures had expanded to 9% and remained the dominant reactivity **(Fig. 2C)**. In PSMB9-targeted cultures, TCRvβ5.1^⁺^cells had expanded to 11%, and around half were detectably reactive. Functional specificity was demonstrated by responses to autologous but not allogeneic tumor organoids **(Fig. 2C)**.

We also expanded day 14 cells that had received neoantigen stimulation but not bead selection (i.e. NeoExpand-REP) and found the frequency of cells expressing the relevant vβ (vβ7.1 or vβ5.1) and that were reactive was substantially lower than for NeoSelect **(Fig. S4A)**, recapitulating the results of experiments with TCR-T cells **(Figs. S3A, S3B)**.

By day 28 we could also detect polyclonal reactivity in multiple vβ families **(Fig. 2D**, **2E)**, especially those expressed by known neoTCRs, suggesting NeoSelect amplified physiologically relevant clonotypes. Peptide titration revealed highly specific NeoPBL responses, but their functional avidities varied between TP53 I254T and PSMB9 R111Q **(Fig. 2D**, **2E)**, indicating NeoSelect can amplify NeoPBL across a broad range of avidities.

To benchmark NeoSelect cultures against TIL workflows, we cryopreserved day 28 NeoSelect products – analogous to the 25–30-day requirement for TIL growth. Each NeoSelect culture yielded well over 1×10D cells, exceeding the ∼1×10D cells typically recovered from TIL fragment cultures prior to REP. Based on *in vitro* assays, we estimated 5–10% of CD8^⁺^ T cells in both day 28 cultures were reactive to TP53 or PSMB9 **(Fig. S4B)**, indicating meaningful enrichment of NeoPBL.

To further quantify NeoPBL frequency, specificity and phenotype within the NeoSelect cultures, we generated an HLA-B*07:02-restricted tetramer that stained positive in 4.9% of CD8^⁺^ T cells in TP53-targeted cultures **(Fig. 2F)**. Phenotypic analysis of tetramer^⁺^ cells revealed 10–19% were CD39^-^ **(Fig. 2G)**, a population rare within CD8^+^ *ex vivo* cultured TIL infusion products administered to patients with cancer. We also observed that some tetramer-positive NeoPBL had high expression of CD28 and/or CD27 **(Fig. 2G)**.

Since many TCRvβ5.1^+^ cells in PSMB9-targeted cultures were reactive, we used this to imperfectly approximate their phenotype in the absence of a tetramer. This showed that vβ5.1^+^ cells also had higher expression of CD28 and/or CD27 compared with non-reactive vβ families **(Fig. 2H, S4C)**. CD28 is a key costimulatory receptor usually absent from neoantigen-specific TIL that is associated with T cell longevity, effector function, and responsiveness to checkpoint blockade, and is a known target of the PD-1/PD-L1 axis^39–41^. These findings indicate NeoPBL exhibit a memory-like, costimulation-competent phenotype, contrasting with conventional GI TIL products^4^.

To gauge NeoSelect’s efficiency, we took day 14 cultures, stimulated them with neoantigen peptides, and then performed magnetic bead extraction. Next, we expanded the bead-captured (‘extracted’) and bead-negative (‘unextracted’) fractions for an additional 14 days **(Fig. S4D)**. By day 28, the extracted pool displayed 160-fold greater neoantigen reactivity than the unextracted pool **(Fig. S3E),** demonstrating that NeoSelect removes many antigen-specific cells.

Additional experiments showed that neither bead selection alone without neoantigen peptide (**Fig. S4F, G)** nor non-specific capture of activated lymphocytes **(Fig. S4H)** accounted for this result. Taken together with the functional assays **(Fig. S4A)**, these results indicate that NeoSelect’s ability to enrich for rare reactive cells depends on true antigen specificity and does not simply amplify high-frequency, cross-reactive T cells, at least in these experiments.

To generate an infusion product, cryopreserved TIL are expanded for an additional 14 days. To model this, we expanded day 28 PBL cultures from patient 4556 with NeoExpand-REP **(Fig. 2I)**. After this, the frequency of vβ7.1^+^ cells had increased to 30% of CD8 T cells in TP53-targeted cultures, and approximately 77% of these were reactive, along with 18% of vβ7.1^⁻^ cells **(Fig. 2J)**. In PSMB9-targeted cultures, the frequency of TCRvβ5.1^⁺^cells had increased to over 50% of CD8 T cells **(Fig. S4I)**. Theoretical yields exceeded 1×10¹² cells each – far more than the maximum tolerated TIL dose.

### Reactivity-seq defines the identity, diversity and transcriptional state of NeoPBL clonotypes

Given the overlap between vβ families expressed by the original neoTCRs and day 28 NeoPBL, we next aimed to determine (i) how many NeoPBL clonotypes there were, (ii) whether they were present in TIL, and (iii) how their transcriptional programs compared to non-reactive PBL that had expanded during 28 days of NeoSelect.

To this end, we analyzed scRNA-scTCRseq by uniform manifold approximation (UMAP), defining clusters based on the transcriptional state of T cells. First, we analyzed TIL from patient 4556. The original neoTCRs recognizing TP53 I254T and PSMB9 R111Q **(Fig. S2B)** had been predicted based on their localization in cluster 9 (TIL_C9), a high scoring ‘neoTCR8’ cluster with gene expression signatures of exhaustion and dysfunction arising from *in vivo* anti-tumor responses and therefore predicted to be highly enriched for neoantigen-specific T cells **(Fig. 3A)**^8^.

**Figure 3.**
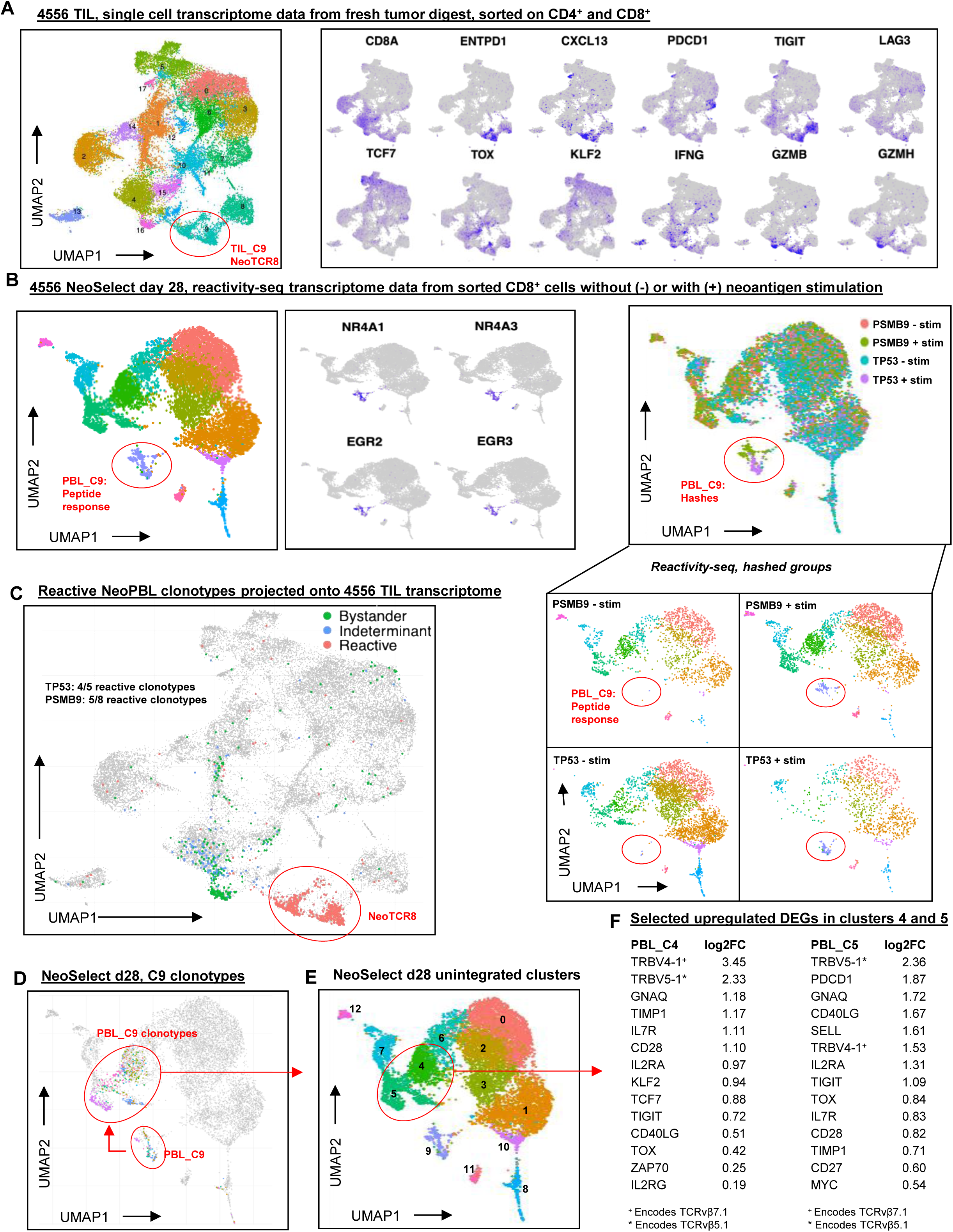
Reactivity-seq defines the identity, diversity, and transcriptional state of NeoPBL. (A) Single-cell transcriptomic data from patient 4556 TIL, showing UMAP clustering; TIL Cluster 9 exhibits the strongest NeoTCR8 signature score and was used to predict the original neoTCRs. (B) Reactivity-seq analysis of day 28 NeoSelect cultures from patient 4556, sorted on CD8⁺ T cells with or without peptide stimulation. Characteristic genes of the reactivity cluster (PBL Cluster 9) are shown in the middle. UMAPs of day 28 NeoSelect cultures combined and separated by hashed sample identifiers are shown on the right. (C) Projection of reactive (confirmed and high-probability) clonotypes onto the 4556 TIL UMAP reveals 9 of 13 NeoPBL clonotypes were present in TIL. The ‘neoTCR8’ cluster is circled in red. Also projected are the bystander (abundant in both NeoSelect cultures, no compelling evidence of reactivity) and indeterminant clonotypes (enriched in one culture, no compelling evidence of reactivity). (D) Reactivity cluster (PBL C9) clonotypes highlighted in resting-state cells from day 28 NeoSelect cultures. (E) Single-cell UMAP of day 28 NeoPBL cultures with annotated clusters. (F) Differential expression analysis comparing upregulated genes in clusters 4 and 5, which were enriched for NeoPBL clonotypes.

To unbiasedly identify neoantigen-specific (NeoPBL) clonotypes that may have been amplified by NeoSelect, we developed a modified scRNA-seq assay, reactivity-seq, which incorporates neoantigen-specific peptide stimulation prior to sequencing, and is therefore able to elicit a distinct transcriptional profile that indicates reactivity. We validated reactivity-seq in 4610 TIL (its neoantigen is described in **Fig. S2A-C)**, detecting the CD8^⁺^vβ17^+^ neoantigen-specific response from a pool of 96 predicted neoantigen peptides **(Fig. S5A-H)**. Although not all reactive clones respond to stimulation in the assay **(Fig. S5F)**, there was high specificity allowing confident identification of neoantigen-responsive clonotypes.

We next applied reactivity-seq to 4556 PBL cultures on day 28, after two NeoSelect cycles, to identify neoantigen-specific clones that may match those discovered in the metastatic tumor deposits or potentially represent new reactivities. The day 28 NeoSelect PBL cluster 9 (PBL_C9) exhibited the peptide response gene expression signature (*NR4A*, *EGR*, *TNFRSF9*, *IFNG*, *TNF*, etc) indicating TCR activation, and was almost exclusively found in neoantigen-stimulated conditions **(Fig. 3B, S6A, S6B)**.

Several clonotypes within PBL_C9 matched the original neoTCRs predicted from TIL_C9, and there were also new putative NeoPBL reactivities. We also identified other high-probability NeoPBL clonotypes through 4-1BB upregulation in response to neoantigen peptide and culture-specific enrichment (i.e. high relative abundance in either the TP53-targeted or PSMB9-targeted NeoSelect culture).

Bystander clones stood out because they were relatively abundant in both the TP53- and PSMB9-targeted NeoSelect cultures and were largely absent from the 4-1BB^+^ fraction and the reactivity-seq (PBL_C9) dataset. The remaining clonotypes were classified as indeterminate – they were relatively enriched in only one of the NeoSelect cultures, but lacked evidence of reactivity by either 4-1BB sorting or reactivity-seq.

In total, we identified 13 NeoPBL clonotypes – 5 TP53-specific and 8 PSMB9-specific. Two of the TP53 reactivities were previously confirmed as neoTCRs whereas 3 were new high probability neoPBL candidates **(Fig. S6A, S6B)**. Independently, the 3 high-probability TP53-specific NeoPBL candidates were also identified by tetramer positive sorting and sequencing. Four of the PSMB9 NeoPBL were previously confirmed neoTCRs and 4 were new high-probability candidates **(Fig. S6A, S6B)**.

Of the 13 NeoPBL clonotypes for both neoantigens, nine were detected in the TIL single-cell atlas **(Figs. S6F, S6G)**. All of the NeoPBL clonotypes amplified by NeoSelect tightly localized to TIL_C9 within the tumor, consistent with the notion that NeoSelect enriches for bona fide anti-tumor activity associated clonotypes *in vivo* **(Fig. 3C)**. Indeterminant PBL clones mapped mostly to TIL_C4 **(Fig. S6C)**. Interestingly, bystander PBL mapped heavily to TIL_C16 **(Fig. 3A, 3C, S6C)**, composed of functional CD8^⁺^ T cells lacking tumor-reactivity signatures. A feature of the physiology of bystander PBL, potentially a proliferative phenotype, led them to transcriptomically cluster *in vivo* (in TIL_C16) and facilitated their expansion *ex vivo* (during NeoSelect).

Finally, we asked whether NeoPBL could be distinguished from bystander PBL based on gene expression after NeoSelect (i.e., on day 28). Analyzing unstimulated cells from reactivity-seq, we saw NeoPBL localized mainly to PBL_C4 and PBL_C5 **(Fig. 3D**, **3E)**. These clusters were characterized by high expression of progenitor- and memory-associated genes (*IL7R*, *TCF7*, *KLF2*, *MYC*) and costimulatory molecules (*CD28*, *CD27*, *CD40LG*) **(Fig. 3F, S6D–E)**. They also expressed exhaustion markers (*TOX*, *PDCD1*, *TIGIT*), consistent with a T_PEX_ phenotype and indicating a T cell population capable of sustaining effector function and proliferative potential^42^. These results are consistent with the notion that circulating anti-tumor T cells are in a less exhausted phenotypic state^22^, and that these pre-exhausted cells can be amplified to the numbers and frequencies often employed in ACT.

### NeoSelect reveals a broad, functional repertoire of tumor-specific T cells targeting unique neoantigens

To fully assess the breadth of reactivity after NeoSelect, we quantified all clonotypes from day 28 cultures. Confirmed or high-probability (reactive) NeoPBL were 7.7% of CD8^⁺^T cells in TP53-targeted but were 27.9% of CD8^+^ T cells in PSMB9-targeted cultures **(Fig. 4A–C)**. Our *in vitro* function-based assays had underestimated PSMB9 reactivity, perhaps due to its lower avidity **(Fig. 2E)**. While most PBL clones were identifiable as bystanders, there were dozens of indeterminant clonotypes – and some localized strongly to PBL_C4/C5 alongside NeoPBL. This suggests transcriptional profiling could infer reactivity after NeoSelect and reveals broad clonotypic diversity.

**Figure 4.**
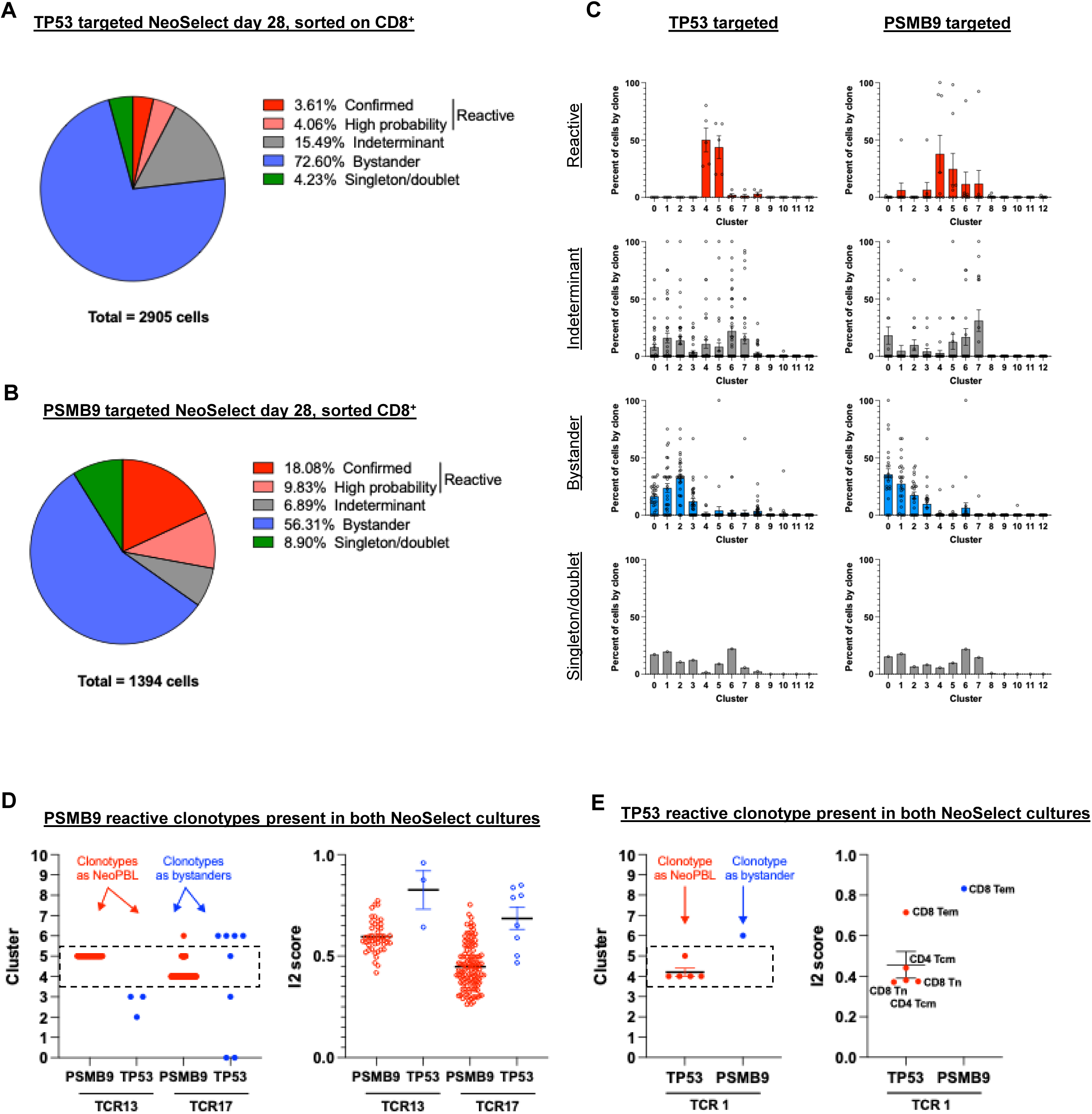
NeoSelect uncovers a broad and functionally poised repertoire of tumor-reactive clonotypes. (A) Pie chart showing the distribution of clonotypes in TP53-targeted NeoSelect cultures categorized as confirmed (cloned and tested), high-probability (based on > 100-fold relative enrichment compared to PSMB9 cultures <u>and</u> evidence of reactivity from 4-1BB sorting or reactivity-seq), indeterminant (> 100-fold relative enrichment but no other supporting evidence of reactivity), or bystander (present in both NeoSelect cultures at similar frequencies). (B) Pie chart showing clonotype classification in PSMB9-targeted NeoSelect cultures using the same criteria as in (A). (C) Single-cell transcriptomic cluster assignment for each TCR identified in TP53-targeted (first column) or PSMB9-targeted (second column) NeoSelect cultures on day 28. Graphs show reactive (red, first row), indeterminant (grey, second row), bystander (blue, third row) and singleton/doublets (fourth row). Each dot shows the percentage of a given clonotype present in the indicated transcriptomic cluster shown in Fig. 3E. (D) Cluster identity and I2 cell type scores for TCR13 and TCR17 when found as reactive NeoPBL (in PSMB9-targeted cultures) or as bystanders (in TP53-targeted cultures). (E) Cluster identity and I2 scores for TCR1 when found as reactive NeoPBL (in TP53-targeted cultures) or as a bystander (in PSMB9-targeted cultures).

To test how neoantigen exposure shapes NeoPBL during NeoSelect, we evaluated three clonotypes (13, 17, and 1) detected in both cultures. Clonotypes 13 and 17, PSMB9-reactive, localized to PBL_C4/C5 in PSMB9-targeted cultures but not in the context of TP53-targeted NeoSelect, where they would have lacked neoantigen stimulation and therefore were effectively bystander PBL **(Fig. 4D)**. Clonotype 1 (TP53-specific) followed the opposite pattern **(Fig. 4E)**. Thus, NeoSelect expands tumor-reactive T cells maintaining at least some clones in a functional, stem-like transcriptional state as compared to bystander PBL.

In addition to patients 4556 (**Fig. 2**, **3**) and 4633 **(Fig. S4G)**, we also tested NeoSelect using samples from patient 4617, an individual also with colorectal cancer. First, we found that low-frequency neoantigen-specific TIL could be enriched by NeoSelect, too **(Fig. S7A-C)**. Next, we expanded PBL with two NeoSelect cycles, after which ∼32% of CD8 T cells were reactive to a CAPZA2 neoantigen and ∼7% of CD8 T cells were reactive to a HAUS4 neoantigen – reproducing the high-level NeoPBL enrichments seen in patient 4556 and 4633 **(Fig. S7D, S7F)**. Also, three reactive TCRvβ families seen in TIL (vβ5.3^⁺^, vβ13.6^⁺^, vβ3^⁺^) were also highly reactive as NeoPBL **(Fig. S7E, S7G)**. TCR sequencing revealed top clonotypes were shared between TIL and NeoPBL among these vβ families, demonstrating NeoSelect faithfully amplifies tumor-relevant clonotypes **(Fig. S7H, S7I)**.

Altogether, reactivity-seq enabled high-resolution mapping of NeoPBL diversity and showed NeoSelect expanded NeoPBL to clinically relevant frequencies. In patient 4556, NeoSelect yielded 13 reactive CD8^⁺^clonotypes and 9 were detectable by scRNAseq in fresh TIL – but none of these had successfully grown during GMP *ex vivo* TIL culture. Importantly, through two cycles of NeoSelect, NeoPBL retained transcriptional evidence of stemness, costimulation, and long-term fitness – signatures often absent from neoantigen-specific GI TIL. These findings position NeoSelect as not only an enrichment tool and TCR-identification strategy, but as a platform for generating a polyclonal, scalable, and potentially effective T cell therapy for patients with solid tumors.

## Discussion

This study presents a new framework for expanding NeoPBL to clinically relevant scales. Conceptually, it addresses two long standing barriers limiting the clinical translation of PBL-derived tumor-reactive T cells: their extremely low frequency and the critical need to preserve their stem-like characteristics. Here we show NeoExpand-REP and NeoSelect – two related culture platforms – can reliably amplify T cells targeting specific neoantigens while preserving critical memory-like phenotypes and potentially restraining differentiation.

NeoExpand-REP not only efficiently expands TCR-T cells but also appears capable of recruiting diverse clonotypic variants and preserving stem-associated markers (e.g., CD39^-^). Consequently, these cells exhibit superior functionality on a per-cell basis *in vitro* and demonstrate efficacy *in vivo*. Importantly, when applying NeoExpand-REP to TIL populations, even cultures initially dominated by highly differentiated cells can undergo potent, reproducible expansion, likely reflecting a selective enrichment of less differentiated subsets rather than direct cellular reprogramming **(Fig. S2)**.

Building on these insights, we developed NeoSelect, a protocol that magnifies the enrichment effect of NeoExpand-REP through bead extraction of T cell-APC conjugates. By performing two NeoSelect cycles in series, its already potent enrichment factor is multiplicative. This enables, over a 28-day period, enrichment of NeoPBL to frequencies comparable to TIL, but from an apheresis, if the targeted antigens are known. Single-cell transcriptomic analyses showed some NeoPBL clonotypes are found in TIL, but retain high expression of genes such as *TCF7*, *KLF2*, *CD28*, consistent with previous reports of pre-exhausted anti-tumor T cells in the circulation of patients with metastatic cancer^22^.

It is also worth emphasizing that these stemness genes (*TCF7*, *CD28*, etc) were upregulated in NeoPBL relative to bystander PBL. It would be more relevant to compare the gene expression of NeoPBL with *ex vivo* expanded TIL on a per clonotype basis, but this was not possible for patient 4556 because CD8^+^ neoantigen-specific TIL did not expand in culture. Nonetheless, these results indicate NeoSelect may yield highly enriched, polyclonal, and phenotypically “younger” T cell pools that could compare favorably to conventional TIL or TCR-T cell products.

NeoSelect was enabled by two core discoveries. The first is that intense on target stimulation, as achieved through both NeoExpand-REP and NeoSelect, does not cause an insurmountable level of AICD and maintains more clonal diversity compared to a REP. From the perspective of an individual T cell, the protocols are virtually indistinguishable, save for 5 minutes during bead extraction. We speculate that the strong initial 16-hour neoantigen stimulation (as shown in **Figs. 1A, S1A, S2E)** selects for memory T cells capable of improved expansion, while perhaps eliminating terminally differentiated cells.

It is known that clonotypes can exist in multiple epigenetic states, and ‘younger’ phenotypes tend to have greater proliferative capacity, so it is possible that NeoSelect acts on those cells. We therefore suggest that cell-intrinsic pathways – exemplified by genes like *KLF2*, *IL7R*, and *TCF7* – may be particularly important targets for future investigations, as they underlie the proliferative and self-renewing capacities of expanded cells.

The second discovery was that we could use magnetic selection to indirectly extract NeoPBL from culture by removing neoantigen-pulsed target cells. This process is not perfectly efficient, because target cell extraction also carries along bystander cells. However, that NeoPBL are disproportionately more likely to be carried with target cells is a key advantage we exploited that allowed NeoPBL to be highly enriched in just 28 days. TCR-peptide-MHC interactions have very weak binding affinity^43^, but in NeoSelect, the sheer number of interactions with target cells pulsed with minimal neoantigen peptides appears to be sufficient to withstand the physical forces involved with magnetic selection.

Our work could provide valuable addition to the developing field of autologous cell therapy for GI cancers. A chief limitation of TIL therapy for these cancers is the requirement for surgical tumor procurement – which is difficult and clinically challenging, but potentially obviated by NeoPBL. The tendency of GI TIL to be highly differentiated and dysfunctional imposes a further barrier. That NeoExpand-REP can efficiently expand highly differentiated TIL is promising **(Fig. S7C)**, but if NeoPBL retain superior phenotypic and transcriptional plasticity, a greater benefit may come from using blood from the start, if the targeted neoantigens are known.

There are two additional potential biological advantages of NeoSelect, although both require additional investigation. First, NeoSelect could theoretically facilitate the targeting of more unique tumor neoantigens. GI malignancies harbor considerable intratumoral heterogeneity, leading to diverse mutational landscapes. Cell therapies directed against a single target neoantigen leave well-trodden paths of tumor escape via antigen or HLA loss. Clinically, most patients who responded to GI cancer TIL in a recent phase II study had infusion products recognizing multiple neoantigens^4^. The key question is how reliably NeoSelect can amplify NeoPBL for a given neoantigen, but answering this will require systematically evaluating *ex vivo* TIL culture and NeoSelect in many patients.

A second potential biological advantage of NeoPBL is that it may enable polyclonal T cell responses against each targeted neoantigen. Here, we demonstrate NeoSelect raised a polyclonal pool of T cells targeting the TP53 and PSMB9 neoantigens from patient 4556, but polyclonality was also evident in the reactivity to CAPZA2 and HAUS4 neoantigens in patients 4617 **(Fig. S7)**. As a point of contrast, individualized TCR-T cell therapies often rely on a monoclonal TCR response, and designing such a treatment may depend on choosing the “right” TCR in the absence of any reliable metric to determine such^3^. And, as we observed in patient 4610, neoantigen-specific TIL responses can be monoclonal, although it is not clear how often this is the case **(Figs. S2 and S5)**. Nonetheless, there could be a theoretical advantage to mounting a polyclonal response against each neoantigen, as it could result in synergistic or complementary killing mechanisms^13^.

Previous studies have employed circulating blood cells to discover anti-tumor TCRs that were administered as TCR-engineered cell therapy^44^ or directly expanded autologous PBL-derived T cells as cell therapy^21^, with minimal durable anti-tumor responses. Because these prior studies did not demonstrate that the administered cells were tumor relevant, it is unclear if the lack of tumor regression were due to lack of tumor-relevant T cells infused.

It is noteworthy to consider the present approach being augmented by checkpoint blockade, by use of costimulatory approaches, and even combination therapy. We saw expanded NeoPBL have moderate-to-high expression of checkpoint molecules like PD-1 and TIGIT, which is also the case for NeoPBL *in vivo*^22^, suggesting they could benefit from PD-1 blockade, TIGIT inhibition, or perhaps 4-1BB agonism^45^. Synergy with checkpoint inhibitors could extend the durability and magnitude of responses *in vivo*. Future studies might test these combinations systematically, assessing how *ex vivo* protocols interact with *in vivo* regimens.

An important limitation of NeoSelect is that it requires *a priori* knowledge of unique patient neoantigens. However, neoantigen discovery can sometimes be performed using primary clinical specimens or even core biopsies^46,47^ and does not need to be performed in GMP facilities. It is also plausible that NeoSelect itself could be used for neoantigen discovery, assuming tumor sequencing is available, which would further broaden its applicability. Finally, it is possible that NeoSelect could be used to raise NeoPBL in a neoantigen-agnostic fashion using pools of predicted peptides, and so could be useful as a simple TCR-discovery tool.

The present work leaves several relevant questions unanswered and clearly indicates the need for additional investigation moving forward. This includes the long-term persistence and *in vivo* efficacy of NeoPBL. While we show potent anti-tumor killing *in vitro* and promising results in mouse models, large-scale clinical validation is needed to evaluate whether NeoPBL can achieve responses in GI cancers or other solid tumors. On a more practical level, it is not clear whether NeoPBL will persist *in vivo* – and whether (or more accurately, how quickly) exhaustion occurs. There remains substantial room for optimizing culture conditions to further enrich CD39^-^ memory cells. This may include identifying the cytokines, small molecules, or additional costimulatory signals that maintain T cell trajectories that preserve memory cells while optimizing enrichment.

Overall, this work may broaden the therapeutic landscape by proposing a simplified route to generating polyclonal, stem-like T cells from peripheral blood using *a priori* knowledge of MHC class I-restricted neoantigens. This approach could enable adoptive cell therapies across many solid tumors and might especially be useful in cancers where disease progression is rapid and/or when harvest of metastatic lesions is particularly burdensome. The demonstration that NeoPBL clonotypes overlap with TIL – yet preserve favorable phenotypic and transcriptional profiles – hints that they may capture the best of both worlds: tumor specificity and robust proliferative potential. The key question is whether NeoPBL can mediate tumor regression in patients; if they can, they could influence how we harness T cells in the fight against cancer.

## Materials and Methods

### Clinical Protocols

All patients were enrolled on NCT00068003 to be screened for NCI Surgery Branch Treatment Protocols, as well as NCT00001823 to collect apheresis and perform tumor harvests, both approved by the IRB of the National Cancer Institute, NIH. Informed consent was obtained and documented in accordance with the Declaration of Helsinki. Patients were 18 years of age or older, ECOG performance status 0 or 1 and were free of systemic infections and major illnesses/comorbidities.

### Patient Tumor Tissue

Patient tissue was obtained from surgical resection of metastatic lesions after discontinuation of systemic therapy for at least one month. Metastases were resected from lung, liver or other sites. Patients had previously undergone a wide range of prior therapies appropriate for their histology and stage such as surgery, chemotherapy, radiotherapy, and/or immunotherapy. The gastrointestinal tumors of patient specimens evaluated in this study were all microsatellite-stable and non-hypermutated.

### Tumor Digestion and TIL sorting for sequencing

Metastatic tumor samples were processed as previously described^1^ using mechanical separation, enzymatic digestion, and gentleMACS dissociation technology (Miltenyi Biotec). Single-cell suspensions of processed tumors were analyzed fresh after resting overnight at 37C in 50/50 media without IL-2.

Lymphocyte-dense fractions of tumor digest were labeled with DAPI (BioLegend) or other viability dyes and stained with fluorescently conjugated antibodies. Viable CD4^+^ or CD8^+^ cells were isolated using an SH800S multi-application cell sorter (SONY Biotechnology) prior to loading sample onto 10x Chromium Controller (10x Genomics).

### Generation and testing of TILs

Fresh tumors were cut into multiple 2-3 mm fragments and cultured in media with cytokine support as described elsewhere^2^. Tumor-specific mutations were identified based on tumor exome and tumor RNA sequencing^2^. TIL reactivity to expressed mutations was identified with peptide pools and tandem mini-genes (TMGs) encoding the products of expressed somatic mutations^3^.

### Flow cytometry antibodies

The following is a list of commonly used antibodies (all from BD Biosciences):

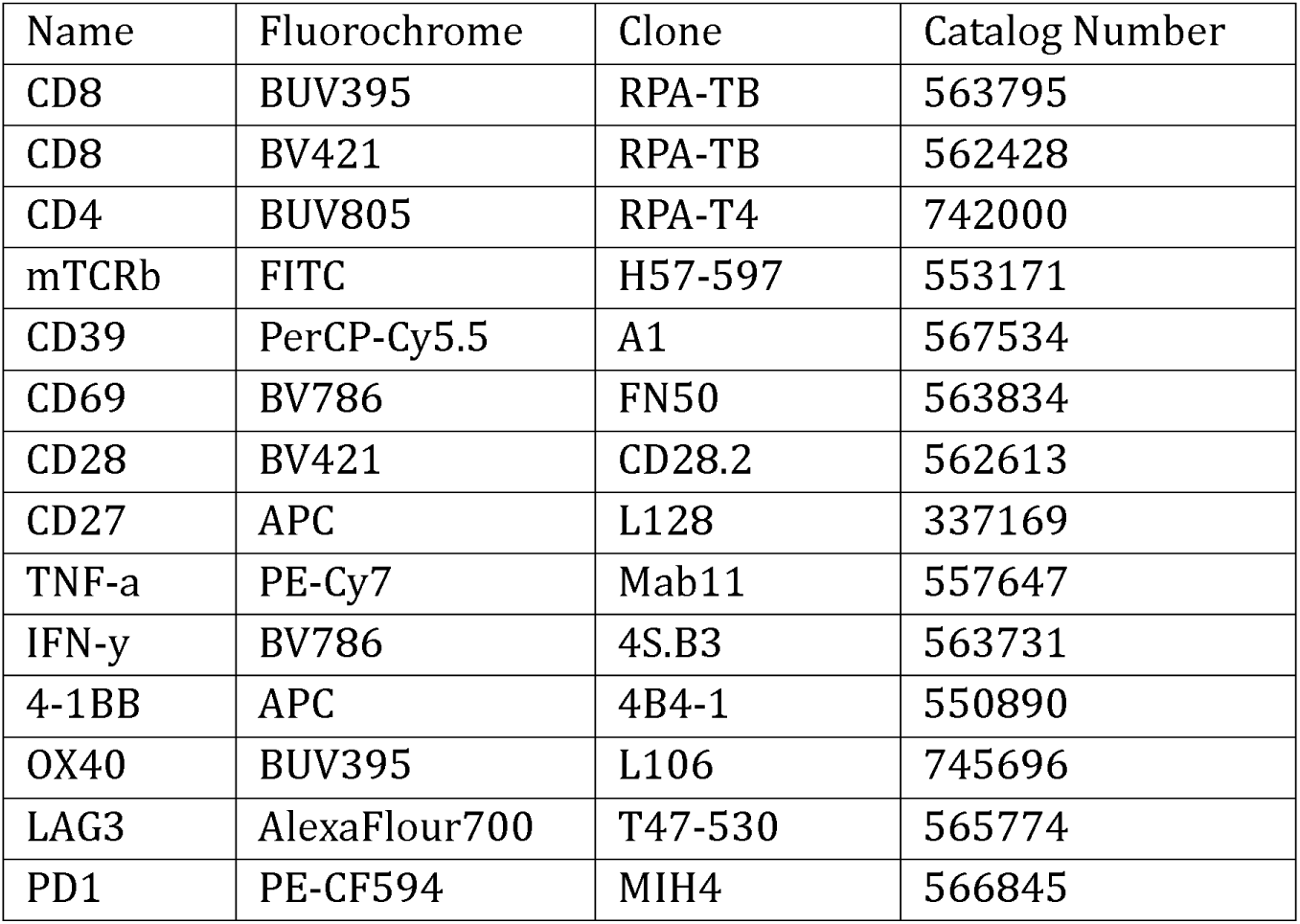

| Name | Fluorochrome | Clone | Catalog Number |
| --- | --- | --- | --- |
| CD8 | BUV395 | RPA-TB | 563795 |
| CD8 | BV421 | RPA-TB | 562428 |
| CD4 | BUV805 | RPA-T4 | 742000 |
| mTCRb | FITC | H57-597 | 553171 |
| CD39 | PerCP-Cy5.5 | A1 | 567534 |
| CD69 | BV786 | FN50 | 563834 |
| CD28 | BV421 | CD28.2 | 562613 |
| CD27 | APC | L128 | 337169 |
| TNF-a | PE-Cy7 | Mab11 | 557647 |
| IFN-γ | BV786 | 4S.B3 | 563731 |
| 4-1BB | APC | 4B4-1 | 550890 |
| OX40 | BUV395 | L106 | 745696 |
| LAG3 | AlexaFluor700 | T47-530 | 565774 |
| PD1 | PE-CF594 | MIH4 | 566845 |

### TCRvβ identification and characterization

TCR vβ tracking was performed via flow cytometry <u>(Beta Mark TCR Vbeta Repertoire Kit, Beckman</u> <u>Coulter)</u> which uses both FITC and PE channels. Twenty-four vβ family surveys for reactivity were performed by doing brief peptide stimulation (see below) followed by individual staining with each of 8 tubes containing 3 unique vβ families – see Beckman Coulter product insert.

### Cell expansion

#### REP

The rapid expansion protocol (REP) was performed as previously described^2,4,5^. Briefly, effector cells (TIL, TCR-T or PBL) are added in a 1:100 ratio with mixed irradiated allogeneic feeder cells plus OKT3 (anti-CD3 antibody, 30 ng/mL, Miltenyi Biotec) and 3,000 IU/mL IL-2 (Aldesleukin) in 50/50 media. On day 5 media is changed; on day 7 cells are split with AIM-V supplemented with IL-2 and replated at approximately 1e6/mL cell concentration; media change and splitting thereafter is performed as needed. Expansions were performed in G-REX 6 well plates (10cm^2^) starting with 5e5 effector cells and 5e7 feeders (irradiated allogeneic apheresis samples pooled from 3 separate donors) except for clinical-scale expansion described in Figure 1, which used G-REX 100 (100 cm^2^) bioreactors and was scaled up to 5e6 effector cells plus 5e8 feeders.

#### NeoExpand-REP

NeoExpand-REP is performed as two distinct steps: neoantigen peptide stimulation for 16 hours and then a REP as above. NeoExpand-REP is defined by a physiologic parameter – reactive cells should be highly activated after the 16-hour peptide stimulation before transfer to REP conditions. This should be quantified by analyzing an aliquot of cells via flow cytometry after the stimulation to measure 4-1BB expression (gating on CD8^+^TCR^+^ or CD8^+^vβ^+^ if known). At least 50% and ideally 70-90% of potentially reactive cells should be 4-1BB^+^ following on-target stimulation (see **Figs. 1A, S1A, S2E)**.

For the peptide stimulation we use as target cells irradiated autologous apheresis or autologous CD4 T cells pulsed with 1 μg/mL minimal neoantigen peptides followed by three washes. In general, a 1:5 E:T ratio (i.e. 5e5 effectors plus 2.5e7 target cells) in a single well of a 24 well plate overnight is sufficient. The stimulation can be titrated as needed to achieve the requisite activation (measured by 4-1BB expression described above) by adjusting the number of target cells, the peptide concentration or even by using a vessel with a smaller surface area (which increases activation by ensuring close contact between effectors & targets).

After antigen-specific stimulation, cells are transferred to REP conditions for expansion in a larger vessel. If starting with 5e5 effector cells, peptide stimulation is performed in a 24 well plate and then the contents are transferred to a GREX 6-well plate (with 5e7 feeders, OKT3 and IL2). If starting with 5e6 effector cells, the peptide stimulation should be performed in a larger vessel (T25 flask or 6-well plate) to ensure adequate media is present over the 16-hour co-culture before final transfer to a G-REX 100 (with 5e8 feeders, OKT3 and IL2).

Finally, it should be noted that short neoantigen peptides immediately bind to class I MHC within 5 minutes and do <u>not</u> require processing like normal peptides. Any nucleated cell, including CD8 T cells, can be used as “APCs” after pulsing with short peptides. Removal of excess minimal peptide by careful washing is crucial because otherwise reactive cells will engage in fratricidal killing if excess peptide is abundant in culture medium. This is the same principle behind brief antigen stimulation, which does not require APCs, described below.

#### NeoSelect

NeoSelect is performed similarly to NeoExpand-REP except that target cells are extracted using magnetic beads after the 16-hour antigen-specific stimulation but before transfer to REP conditions. This implies the surface molecule recognized by beads must be present on target cells but not on effector cells. Purified autologous lymphocytes transduced with truncated nerve growth factor receptor (NGFR) work well for this purpose, as they can be extracted using NGFR^+^ microbeads (Stem Cell Tech).

More simply, target cells can simply be purified autologous CD4^+^ T cells and therefore can be extracted using CD4^+^ microbeads (also Stem Cell Tech). NeoSelect starts with a CD8 memory enrichment (Stem Cell Tech) on day 0, and an additional CD8 negative enrichment (Stem Cell Tech) should be performed on day 14 if needed, so CD4^+^ selection works as an alternative to NGFR^+^ selection although the efficiency may be slightly lower.

Also, because bead extraction leaves behind many bystander CD8^+^ cells, NeoSelect can be scaled up relative to NeoExpand-REP. In other words, NeoExpand-REP starts with 5e5 effectors because that is the maximum starting population of a 6-well GREX. NeoSelect can be scaled up (e.g. 5e6 effectors:2.5e7 targets) because only ∼10% of CD8+ effectors (5e5) are extracted during bead selection, if it is efficient. Maintaining a clear biological negative control is critical for quality control during NeoSelect and for subsequent analysis to identify reactive and bystander cells. If using NeoSelect to target two neoantigens, each culture can serve as the other’s negative control (as we did with PBL from 4556). If only one neoantigen is pursued, then the negative control should be extraction of target cells pulsed with DMSO rather than neoantigen peptide (as shown in **Figs. S4F, S4G**).

### Response to overnight stimulation using tumor, organoid or peptide-pulsed target cells

Cytokine responses to cultured <u>TYK-nu</u> tumor cells **(Figures 1F, S1F, S1G)** were evaluated by 16 hour co-culture with sorted CD8^+^mTCRβ^+^ cells using fluorescence-activated cell sorting (FACS) either in 24 well or 96 well plates at titrated E:T ratios ranging from 1:8 to 4:1. After overnight culture, brefeldin and monensin (BD biosciences) was added and an additional 5 hours of co-culture was performed followed by live/dead staining (Thermo-Fisher), fixation/permeabilization (BD biosciences) and intracellular cytokine staining (BD biosciences). Overnight responses to short neoantigen peptide stimulation were obtained by pulsing autologous target cells with short peptides followed by 3 washes and then co-culture with effector cells **(Figs. 2C, S2K, S4A)**.

### Cytokine response to brief neoantigen peptide stimulation

A brief neoantigen stimulation followed by intracellular cytokine staining is used to identify reactive cells with a high degree of specificity. This is accomplished by adding brefeldin plus monensin at indicated amounts (BD biosciences) to the cell pool of interest. Next, minimal neoantigen peptide(s) are added directly to effector cell pools at approximately 1 µg/mL. Minimal peptides rapidly bind to MHC-I molecules (i.e. do not require processing by APCs) and elicit reactivity immediately as reactive cells respond to nearby cells (including other reactive cells) with MHC-bound peptides.

After 3 hours of stimulation, cells are stained for viability (using a fixable dye, ThermoFisher), fixed and permeabilized, and stained for TNF-α, IFN-γ, and any other cytokines or proteins of interest (BD biosciences). Staining of vβ can also be performed on fixed cells. Brief neoantigen stimulation must be performed on a short timescale, as short peptides will elicit fratricidal killing among reactive populations. As shown in **Fig. S4E**, brief neoantigen stimulation is extremely specific: it results in a strong cytokine production by reactive cells (vβ7.1^+^) but virtually no cytokine is detected in bystander cells (vβ3^+^).

### Incucyte assays

To evaluate the cytotoxicity of TCR-T cell products, human ovarian cancer TYK-nu Red cells (5,000 cells/well) expressing red fluorescence protein were seeded into flat-bottom, tissue culture-treated, non-pyrogenic polystyrene 96-well plates. After 1 hour of incubation, TYK-nu were co-cultured with T cells harboring TP53-specific TCRs at titrated effector-to-target (E:T) ratios in 50/50 medium without IL-2. TYK-nu Red cells cultured without T cells served as control. Plates were imaged every 4 hours using an IncuCyte S3 live-cell analysis system (Sartorius, Germany). Cytotoxic activity of the TCR-T cells was assessed by quantifying the Red Image Mean of the TYK-nu Red cells over time using IncuCyte software.

### *In vivo* tumor treatment

NSG mice (NOD-*Prkdc^em26Cd52^Il2rg^em26Cd22^*/NjuCrl, Charles River) were injected subcutaneously with 1e7 TYK-nu cells (which natively express TP53 R175H and HLA-A*02:01) which were allowed to establish for 2 weeks. Titrated doses of TCR-T cells were transferred intravenously via tail vein injection along with IL-2 given intravenously for one dose (along with TCR-T cells) followed by intraperitoneal doses of IL-2 (180,000 IU) on days 2 and 3. Tumor sizes (long axis & perpendicular axis) were measured at least weekly by an investigator blinded to treatment allocation. The use of animals for this study was approved by the National Cancer Institute Animal Care and Use Committee.

### Tumor Organoid Establishment and Culture

Fresh tumor tissues obtained from surgical resections were processed to establish organoid cultures following previously published protocols^6^. Briefly, tissues were mechanically dissociated and enzymatically digested using a human tumor dissociation kit (Miltenyi Biotec). The resulting cell suspension was mixed with 80% Matrigel and 20% colorectal cancer-specific medium and plated in tissue culture-treated plates. After Matrigel solidification, warmed colorectal cancer medium was added. Cultures were maintained in a 37°C CO2 incubator and passaged every 1-2 weeks as necessary^6^.

### Organoid Preparation and Co-culture Experiments

Prior to co-culture experiments, established organoid lines were harvested and digested into single cells using TrypLE (Gibco). Cells were then resuspended in colorectal-specific medium and incubated overnight in Costar low-attachment 6-well plates. For the IFN-γ pretreated condition, 10 ng/ml recombinant human IFN-γ (Peprotech) was added to the medium. Following incubation, organoids were harvested, washed three times in 10 ml of Advanced DMEM by centrifugation (1500 rpm for 5 min), and finally resuspended in Advanced DMEM for subsequent experiments. Responses to allogeneic and autologous organoid with and without pretreatment with IFN-γ **(Figs. 2C and S4A)** were performed overnight with viability and 4-1BB staining the following day.

### Single cell RNA sequencing and analysis

Single cell capture, library preparation, sequencing, data processing and analysis was performed as recently described^7^. For cluster-based scRNA analysis, the FindAllMarkers function (using default values) within Seurat was used to identify differentially expressed genes between clusters. Genes with an adjusted p-value of < 0.05 were included as differentially expressed genes (DEGs).

To establish the D50 we sorted clonotypes by abundance/clones (most abundant to least) until their abundances cumulatively reach 50% of the total population. The D50 index is the number of clonotypes needed to hit the 50% mark.

### TCR Clonotype Selection and Differential Gene Expression Analysis

VDJ sequencing data were processed using R (v4.2.0) with Immunarch (v0.6.8) and Seurat (v4.3.0) to identify dominant TCR clonotypes **(Fig. 1K)** and assess gene expression differences between NeoExpand-REP and REP conditions. For each sample, productive TRB clonotypes (is_cell == true) were grouped by CDR3 amino acid sequence, and clonal abundance was calculated based on cell counts. Clonotypes were ranked by proportional representation, and cumulative frequencies (cum_freq) were computed to assess clonal dominance. To account for variation in sequencing depth and repertoire diversity, dominant clonotypes were selected using dataset-specific cumulative frequency thresholds. Shared clonotypes between NeoExpand-REP and REP samples were identified by intersecting the respective top clonotype sets. Single-cell RNA-seq profiles were annotated according to TRB CDR3 matches, classifying cells as either “shared dominant” or “not shared.” Differential gene expression analysis was performed in Seurat (Wilcoxon rank-sum test) between NeoExpand-REP and REP shared dominant clonotypes, using log fold-change > 0.25 and minimum expression in ≥25% of cells as thresholds. This targeted approach enabled transcriptional comparisons focused on clonally expanded and biologically relevant T cell populations while controlling for clonal and technical variability across samples.

### Short peptide prediction

Short peptides were predicted from tumor-specific mutations based on expression (Max Read2Count > 2) and mutant binding percentile rank < 0.5 (via MHCflurry^8^). Short peptides were first tested *in vitro* to evaluate their ability to elicit reactivity (see below).

### Short peptide sequences

For patient 4556, the mutated neoantigen and wild type sequences are as follows:

|  |  |  |  |
| --- | --- | --- | --- |
| • TP53 | p.I254T | Mut: RPILTTITL | Wt: RPILTIITL |
| • PSMB9 | p.R111Q | Mut: QEDLSAHLM | Wt: REDLSAHLM |

For patient 4617, the mutated neoantigen and wild type sequences are as follows:

|  |  |  |  |
| --- | --- | --- | --- |
| • CAPZA2 | p.I155F | Mut: FACIESHQF | Wt: IACIESHQF |
| • HAUS4 | p.R80M | Mut: LMSEILHRV | Wt: LRSEILHRV |

### Reactivity-seq

Reactivity-seq is a streamlined version of our short-duration neoantigen peptide stimulation protocol. It takes advantage of the fact that brief neoantigen peptide stimulation (described above) is extremely specific because it results in minimal to no activation of bystander cells **(Fig. S4E)** and also preserves cell viability. Briefly, cells are incubated for 120 minutes with short peptides plus brefeldin A and monensin as described for brief neoantigen stimulation above. Following stimulation, cells are washed once with FACS buffer (without EDTA), stained on ice for 30 minutes with cell hashing antibodies and surface markers (e.g., CD4, CD8), washed three times, and resuspended in DAPI for viability.

We sort equal numbers of CD8^⁺^ T cells (usually setting a limit at 50,000 per condition) for single-cell capture. CD4^⁺^T cells are excluded by the sort, or can be depleted by beads up front (Stem Cell Technologies), since MHC class II restricted T cells do not reliably respond to minimal peptides and so cannot be picked up by this version of reactivity-seq. Sorted cells are then pooled (assuming they are hashed) and resuspended at 1×10D/mL (e.g., 30,000 cells in 30 µL) for loading onto the single-cell platform.

In **Supplemental Figure 4**, we stimulated CD8-enriched TIL with mutant peptide pools including all highly predicted binders (100 ng/peptide × 96 peptides; ∼10 µg/mL total peptide concentration). Lower-avidity peptides may require higher concentrations or longer stimulation to generate a clear transcriptional “reactivity cluster.” Reactivity-seq conditions can be titrated by simply performing a pilot experiment using intracellular cytokine staining to confirm there is a detectable response (as shown in **Fig. S5B**). We identify neoantigen-responsive clonotypes by transcriptional clustering **(Figs. 3B, S5C)**.

### NGFR-tagged target cells

To generate tagged target cells, we transduced autologous lymphocytes with with truncated nerve growth factor receptor (NGFR) using the same MSGV1 vector as for transgenic TCRs^5^. Briefly, autologous T cells or apheresis were stimulated with CD3/CD28 beads (ThermoFisher) or OKT3 respectively and transduced with the vector on day 2. Next, NGFR transduced cells are expanded for 10 days. After expansion, CD4^+^NGFR^+^ cells can be sorted using CD271 (NGFR) antibodies (BD Biosciences) or isolated by magnetic selection (Stem Cell Tech) by doing a CD4 negative enrichment followed by NGFR positive selection and expanded further as a purified population in a REP. After 14 days of expansion, purified NGFR^+^ cells can be banked in aliquots of 1e8 and cryopreserved for future use.

