## Supplemental Figure 1 for "Rapid enrichment of progenitor exhausted neoantigen-specific CD8 T cells from peripheral blood"

### Supplemental Figure 1. Neoantigen peptide stimulation prior to a Rapid Expansion Protocol (REP) increases blood-derived TCR-T cell expansion and enriches for a less differentiated T cell phenotype

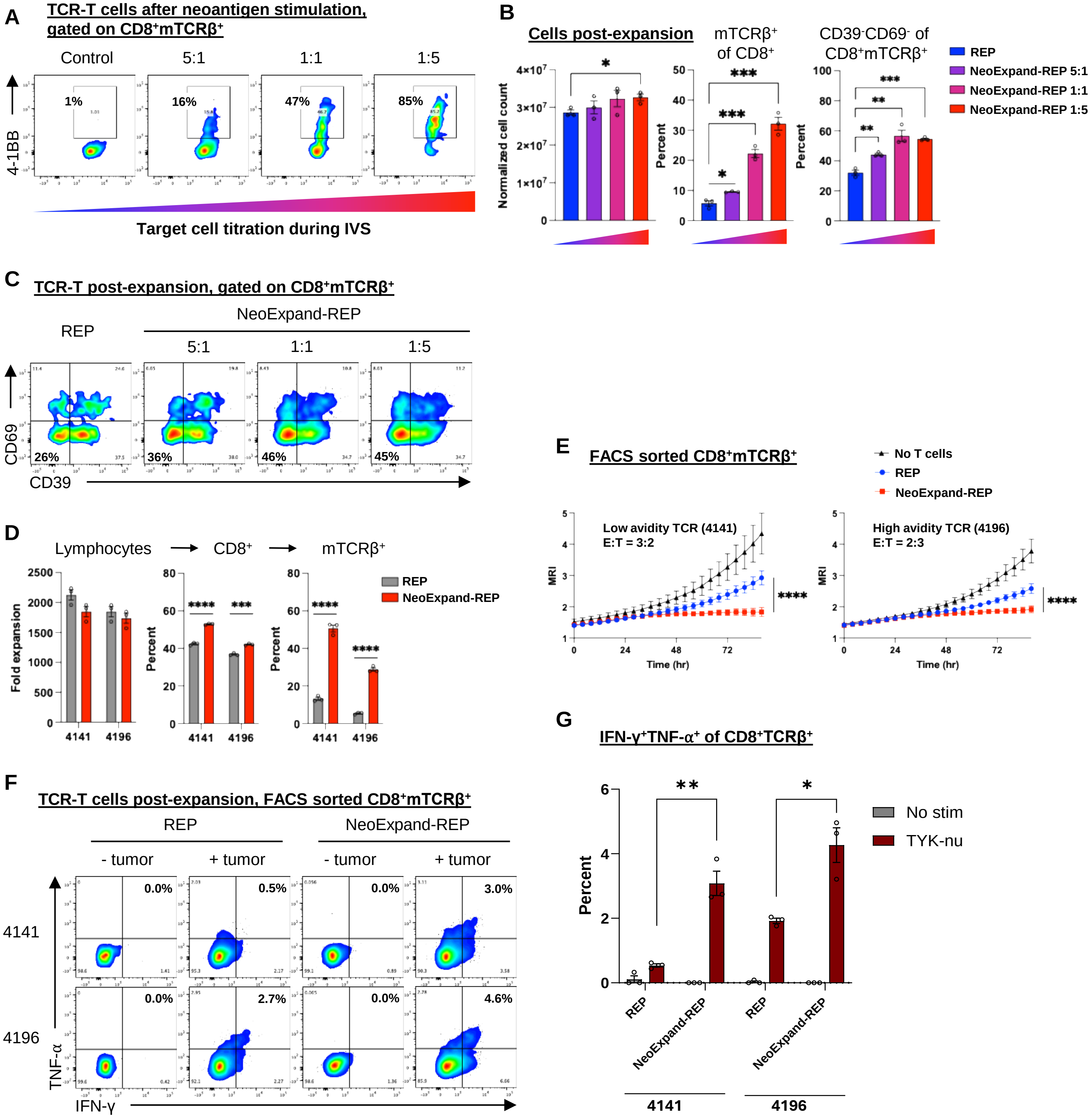

**Supplemental Figure 1. NeoExpand-REP enhances TCR-T cell expansion and function without promoting terminal differentiation.** (A) TCR-T cells targeting KRAS G12D and restricted by HLA-A*11:01 were generated from patient PBL. Flow cytometry plots show 4-1BB expression on $CD8^{+}mTCR\beta^{+}$ cells after 16 hours of neoantigen stimulation, using equal numbers of effector cells with five-fold titrations of irradiated target cells (autologous PBL pulsed with minimal KRAS neoantigen peptide) or control. (B) After neoantigen peptide stimulation, TCR-T cells were transferred to identical REP conditions, expanded for 14 days and then analyzed. Shown are normalized total cell counts, the frequency of $mTCR\beta^{+}$ cells among $CD8^{+}$ T cells, and the proportion of $CD8^{+}mTCR\beta^{+}$ cells expressing the $CD39^{-}CD69^{-}$ phenotype. The effector to target ratio reflects the ratio during the initial neoantigen stimulation. (C) Representative flow cytometry plots showing CD39 and CD69 expression in $CD8^{+}mTCR\beta^{+}$ T cells following REP or NeoExpand-REP expansion. (D) TCR-T cells using receptors isolated from patients 4141 and 4196, targeting TP53 R175H with HLA-A*02:01 restriction, were expanded from patient PBL using REP or NeoExpand-REP. Shown are overall lymphocyte fold expansion (including all $CD4^{+}$ and $CD8^{+}$ T cells), the $CD8^{+}$ fraction of lymphocytes and the $mTCR\beta^{+}$ fraction of $CD8^{+}$ cells. (E) Incucyte assay comparing tumor cell killing by equal numbers of FACS-sorted $CD8^{+}mTCR\beta^{+}$ T cells expanded via REP or NeoExpand-REP. Target cells were TYK-nu human ovarian cancer cells (natively expressing TP53 R175H and HLA-A*02:01). (F) Flow cytometry plots showing intracellular TNF-α and IFN-γ expression in $CD8^{+}mTCR\beta^{+}$ T cells following 16-hour co-culture with TYK-nu cells after REP or NeoExpand-REP expansion. (G) Summary of cytokine co-expression data from (G), showing the frequency of $TNF\text{-}\alpha^{+}IFN\text{-}\gamma^{+}$ double-positive cells gated on $CD8^{+}mTCR\beta^{+}$.
