## Supplemental Figure 2 for "Rapid enrichment of progenitor exhausted neoantigen-specific CD8 T cells from peripheral blood"

### Supplemental Figure 2. NeoExpand-REP enables expansion and functional preservation of neoantigen-specific TIL in an advanced state of differentiation

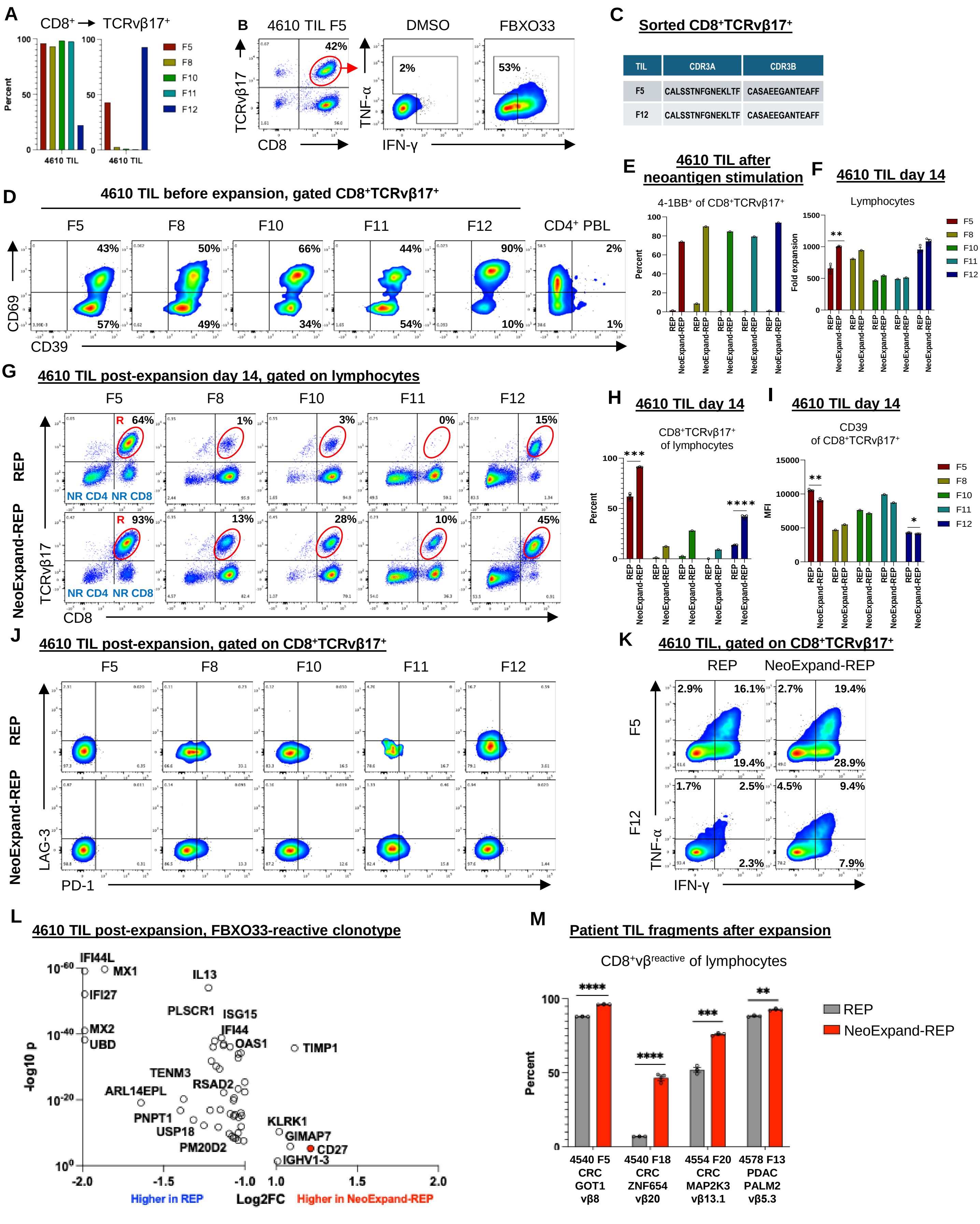

**Supplemental Figure 2. NeoExpand-REP enables expansion and preserves function of terminally differentiated, neoantigen-specific TIL.** (A) The frequency of $CD8^+$ gated on lymphocytes (left) and $v\beta17^+$ gated on $CD8^+$ (right) in five TIL fragments from patient 4610. (B) Flow cytometry data showing CD8 and TCRvβ17 expression on 4610 TIL, and intracellular TNF-α and IFN-γ in $CD8^+TCRv\beta17^+$ cells following stimulation with FBXO33 mutant peptide solubilized with DSMO vs. DSMO alone. (C) CDR3 sequences of $CD8^+TCRv\beta17^+$ cells sorted from 4610 TIL. (D) Flow cytometry data showing CD39 and CD69 expression on $CD8^+TCRv\beta17^+$ TIL at rest, before expansion. (E) Frequency of 4-$1BB^+$ on $CD8^+TCRv\beta17^+$ cells after 16 hours of neoantigen peptide stimulation using autologous peptide-pulsed lymphocytes. (F) Overall lymphocyte fold expansion after 14 days. Technical replicates were only performed for fragments 5 and 12. (G) Representative flow cytometry data showing CD8 and TCRvβ17 expression after expansion with a REP or NeoExpand-REP. R indicates reactive TIL, NR CD4 indicates non-reactive $CD4^+$ TIL, and NR CD8 indicates non-reactive $CD8^+$ TIL. (H) Bar graphs summarizing data from (G). (I) Bar graphs showing CD39 expression on $CD8^+$ $TCRv\beta17^+$ TIL after expansion. Mean fluorescence intensity (MFI) was shown because T cells were universally $CD39^+$. (J) Expression of inhibitory receptors LAG-3 and PD-1 on $CD8^+TCRv\beta17^+$ TIL after expansion. (K) Representative flow cytometry data showing TNF-α and IFN-γ expression of $CD8^+TCRv\beta17^+$ post-expansion TIL after neoantigen peptide stimulation. (L) Volcano plot showing DEGs within the FBXO33 neoantigen-specific TIL clonotype. All genes with log2FC > 1 or < -1 are shown. (M) Bar graphs showing results after 14 days of expansion with REP or NeoExpand-REP using TIL from three different patients recognizing four unique neoantigens. Bar graph shows the percentage of lymphocytes that were positive for CD8 and the unique vβ expressed by neoantigen-specific TIL. The patient ID, TIL fragment culture, cancer type, specific neoantigen and vβ of reactive TIL are indicated. Abbreviation: CRC, colorectal cancer; PDAC, pancreatic ductal adenocarcinoma.
