## Supplemental Figure 3 for "Rapid enrichment of progenitor exhausted neoantigen-specific CD8 T cells from peripheral blood"

### Supplemental Figure 3. NeoSelect enables rapid enrichment of polyclonal, neoantigen-specific CD8$^{+}$ T cells from blood

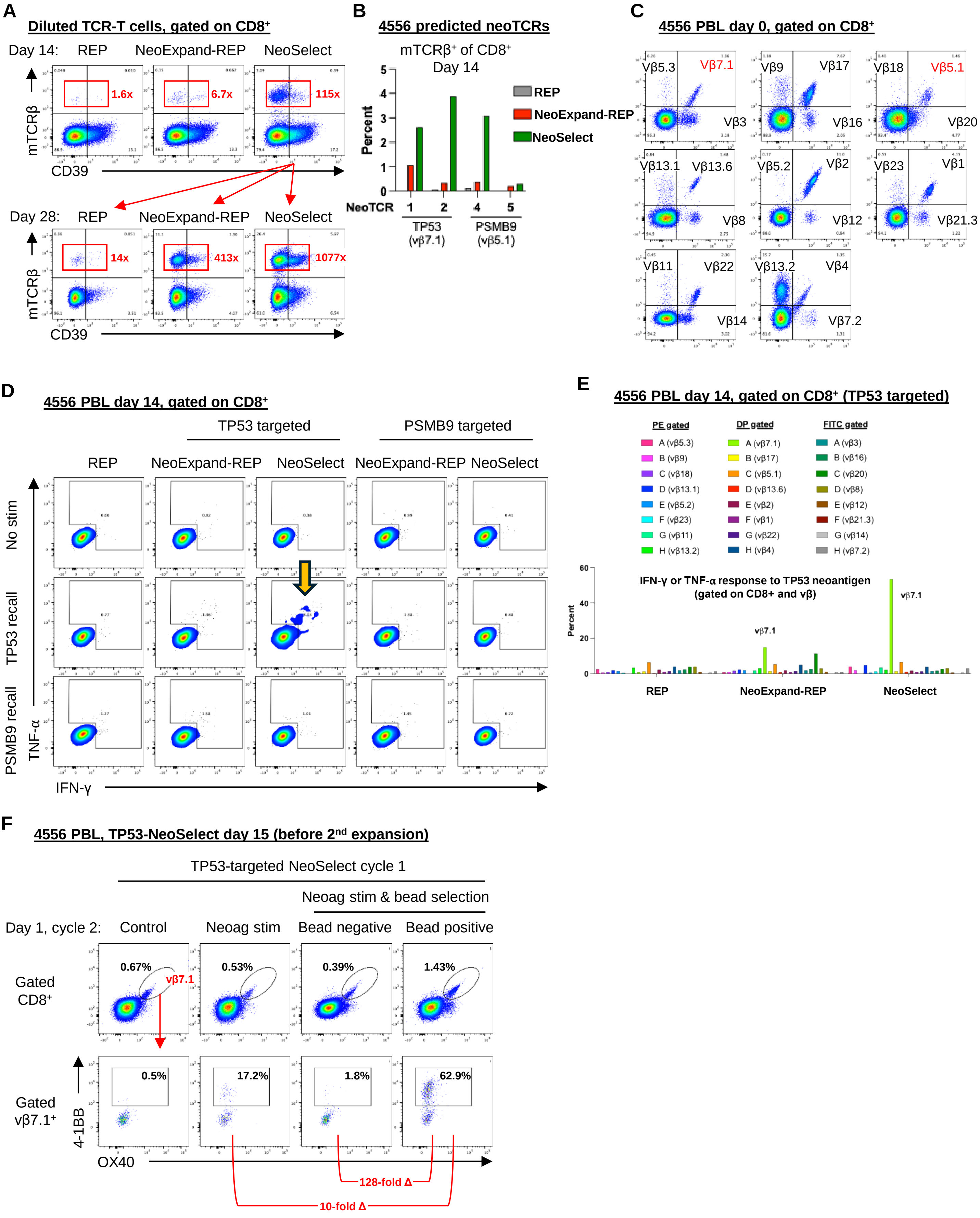

**Supplemental Figure 3. NeoSelect amplifies enrichment of reactive cells during NeoExpand-REP.** (A) Flow cytometry data from a mock NeoSelect experiment using TCR-T cells diluted to approximately 0.03% with untransduced autologous lymphocytes. Plots are gated on $CD8^+$. The top row show the results of an initial 14-day expansion with REP, NeoExpand-REP or NeoSelect, with the relative fold enrichment from the starting frequency shown in red. Bottom row plots show results on day 28 after a second cycle was performed using NeoSelect cycle 1 cells, with the relative fold enrichment from the starting frequency shown in red. (B) Bar graphs showing results from a mock NeoSelect experiment as in (A) using diluted 4556 neoTCRs specific for TP53 or PSMB9 neoantigens. Only one cycle was performed, with bar graphs showing the percentage of $mTCR\beta^+$ gated on $CD8^+$ on day 14 of each condition. (C) TCRvβ repertoire of $CD8^+$ T cells from patient 4556 peripheral blood after $CD8^+$ memory enrichment on day 0. (D) Flow cytometry plots showing cytokine responses of day 14 PBL at rest or following brief stimulation with TP53 or PSMB9 neoantigen peptides. (E) Bar graphs summarizing cytokine responses to brief neoantigen stimulation across 24 common TCRvβ families in $CD8^+$ T cells after 14-day expansion with REP, NeoExpand-REP, or NeoSelect. (F) Flow cytometry plots showing day 15 PBL that were first expanded with one cycle of NeoSelect targeting TP53 over 14 days, and then subjected to 16 hours of IVS with or without bead selection. Top plots show vβ7.1 expression gated on $CD8^+$, while bottom plots show 4-1BB and OX40 expression on $CD8^+v\beta7.1^+$ gated cells. Below the bottom plots, red brackets and numbers indicate the fold difference in $CD8^+v\beta7.1^+4\text{-}1BB^+$ cells between the indicated groups
