## Supplemental Figure 4 for "Rapid enrichment of progenitor exhausted neoantigen-specific CD8 T cells from peripheral blood"

### Supplemental Figure 4. NeoSelect enriches NeoPBL to relevant frequencies.

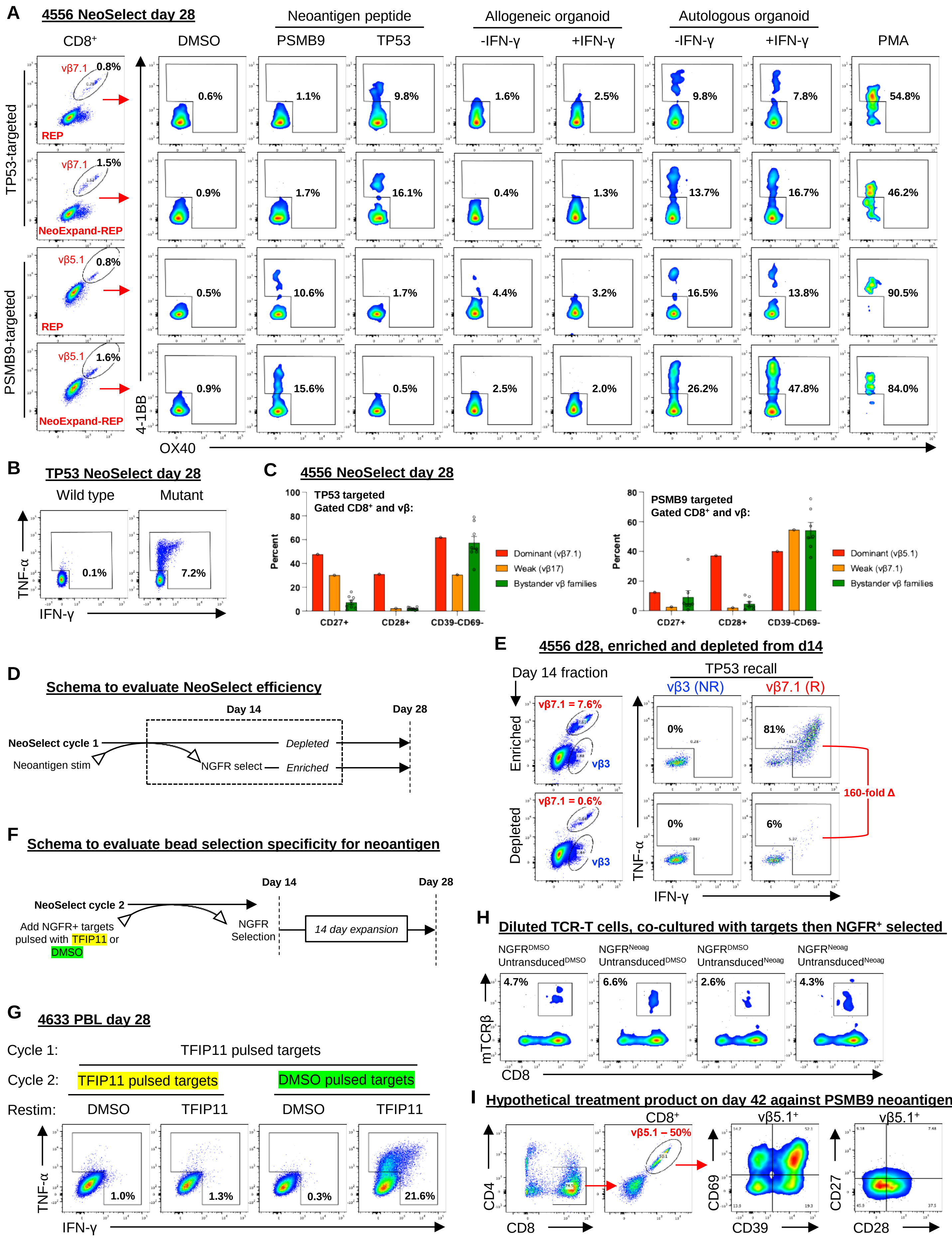

**Supplemental Figure 4. NeoSelect enriches NeoPBL to clinically relevant frequencies.** (A) Day 14 NeoSelect-expanded PBL were put through a second cycle of expansion using a REP or NeoExpand-REP (in parallel with a second cycle of NeoSelect shown in **Fig. 2C**). Flow plots show 4-1BB and OX40 expression on day 28 $CD8^+$ T cells following 16-hour co-culture with neoantigen peptides or tumor organoids. Gated on $v\beta7.1^+$ (TP53) or $v\beta5.1^+$ (PSMB9). (B) Cytokine response of day 28 PBL following brief stimulation with TP53 wild type or mutant peptide. (C) Frequency of $CD27^+$, $CD28^+$, and $CD39^-CD69^-$ $CD8^+$ T cells, stratified by TCRvβ families showing dominant, weak, or no neoantigen reactivity. (D) Schematic of NeoSelect efficiency test: following the first cycle (day 14), bead-enriched (positive) and bead-depleted (negative) fractions were expanded separately. (E) Flow cytometry plots from day 28 cultures showing the frequency of $v\beta7.1^+$ cells and their response to neoantigen peptide. The frequency and reactivity of bystander ($v\beta3^+$) cells are shown for comparison. The red bracket and text indicates the overall fold difference in reactive cell frequency between enriched and depleted cells. (F) Schematic of bead selection specificity test: following the first cycle (day 14), target cells were pulsed with DMSO or neoantigen peptide and continued through a second cycle of NeoSelect. (G) Flow cytometry plots from day 28 cultures showing $CD8^+$ T cells and their response to rechallenge after the test described in (F). (H) Results of an experiment testing the dependence of NeoSelect on neoantigen peptide being present on bead extracted target cells, as opposed to being in culture. Equal numbers of $NGFR^+$ target cells and untransduced target cells were pulsed with DMSO or neoantigen peptide and co-cultured with diluted TCR-T cells before bead extraction. (I) Flow cytometry plots from day 42 cultures from patient 4556 after two cycles of NeoSelect and one cycle of NeoExpand-REP utilizing the PSMB9 neoantigen.
