## Supplemental Figure 5 for "Rapid enrichment of progenitor exhausted neoantigen-specific CD8 T cells from peripheral blood"

### Supplemental Figure 5. Reactivity-seq identifies a single known neoantigen-specific CD8$^+$ clonotype in 4610 TIL.

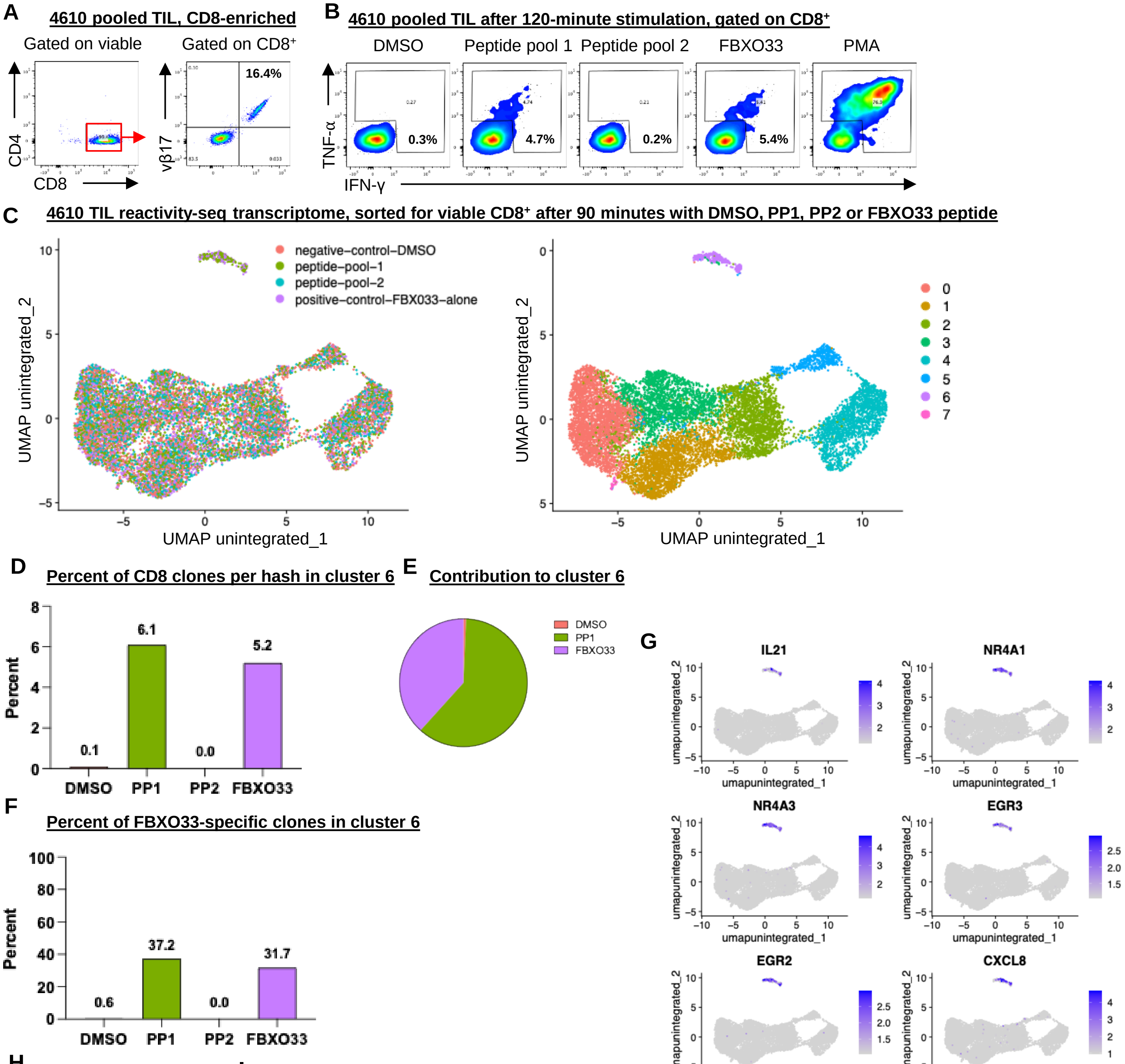


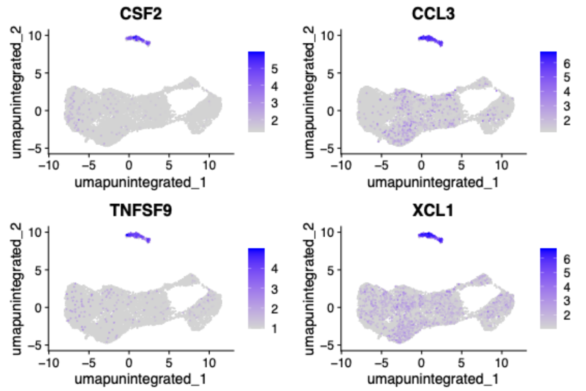


**H** Top upregulated genes

| Gene | log2FC |
|---|---|
| IL21 | 11.2 |
| NR4A1 | 10.2 |
| NR4A3 | 9.9 |
| EGR3 | 9.9 |
| EGR2 | 9.7 |
| CXCL8 | 8.9 |
| CSF2 | 8.6 |
| CCL3 | 8.4 |
| TNFSF9 | 8.2 |
| XCL1 | 8.1 |
| CCL4 | 7.8 |
| CCL3L1 | 7.6 |

**I** Notable upregulated genes

| Gene | log2FC |
|---|---|
| TNF | 7.0 |
| IFNG | 6.2 |
| FASLG | 4.9 |
| CD69 | 4.6 |
| CD40LG | 4.1 |
| TNFSF14 | 4.0 |
| TNFRSF9 | 3.7 |
| PDCD1 | 2.9 |
| TRBV19* | 2.5 |
| CTLA4 | 2.4 |
| ICOS | 2.2 |
| CD109 | 1.9 |

*encodes TCRvβ17

**Supplemental Figure 5. Reactivity-seq identifies a single neoantigen-specific $CD8^+$ clonotype in 4610 TIL.** (A) Flow cytometry plots showing CD4 and CD8 expression in pooled 4610 TIL and the frequency of TCRvβ$17^+$ cells within the $CD8^+$ gated compartment. (B) Cytokine responses following mock reactivity-seq stimulation (120 minutes with peptide and brefeldin/monensin) using DMSO, peptide pool 1, peptide pool 2, FBXO33 alone, or PMA. Each peptide pool contained 96 predicted neoantigen peptides; FBXO33 was in pool 1 and tested as a positive control at the same concentration. (C) UMAP from reactivity-seq transcriptomic data showing $CD8^+$ T cells after stimulation as in (B), excluding PMA-stimulated conditions. The reactivity cluster 6 (C6) appears as a distinct island at the 12:00 position. (D) Proportion of CD8 clones from each hashed condition that were in C6. (E) Pie chart showing the hashed group of all clones found within C6. (F) Percentage of FBXO33-reactive (i.e. $CD8^+$TCRvβ$17^+$) clones that mapped to C6 in each hashed group. (G) UMAPs showing top DEGs in C6. (H) Table of top upregulated genes in C6 with $\log_2$ fold changes. (I) Table of notable DEGs in C6 including TRBV19 (encoding TCRvβ17).
