## Supplemental Figure 6 for "Rapid enrichment of progenitor exhausted neoantigen-specific CD8 T cells from peripheral blood"

### Supplemental Figure 6. Reactivity-seq of day 28 NeoPBL from patient 4556.

**A**

**Reactivity-seq C9 – top DEGs**

| Gene | log2FC |
|---|---|
| IL2 | 10.1 |
| RRAD | 10.1 |
| CD200 | 8.9 |
| NR4A1 | 8.7 |
| EGR2 | 8.4 |
| NR4A3 | 8.3 |
| EGR3 | 8.2 |
| IL13 | 8.1 |
| CCL3L1 | 7.8 |
| CSF2 | 7.6 |
| AC055854.1 | 7.22 |
| PTGS2 | 7.0 |
| EGR1 | 6.8 |

**B**

**Reactivity-seq C9 – notable DEGs**

| Gene | log2FC |
|---|---|
| TNF | 6.7 |
| IFNG | 5.2 |
| TNFSF9 | 4.9 |
| PDCD1 | 4.9 |
| CD40LG | 4.9 |
| TNFRSF9 | 4.8 |
| FASLG | 4.5 |
| CD69 | 4.1 |
| CD109 | 3.9 |
| TRBV6-4* | 3.6 |
| TRBV4-1[+] | 2.1 |
| TRBV5-1[%] | 1.6 |
| TRBV29-1[#] | 0.9 |

*encodes TCRvβ6-4
+encodes TCRvβ7.1
%encodes TCRvβ5.1
#encodes TCRvβ4

**C**

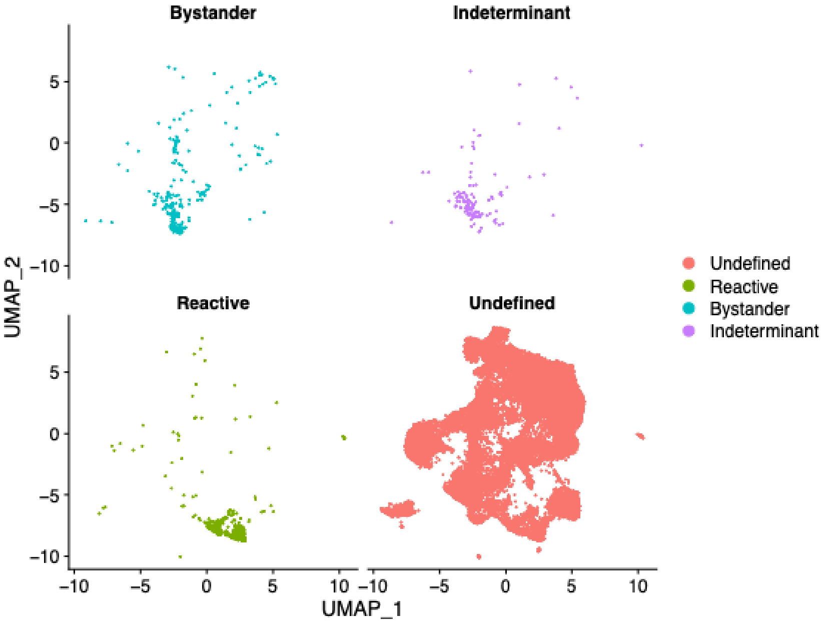


**D**

#### NeoSelect d28 reactivity-seq, selected cluster 4 DEGs

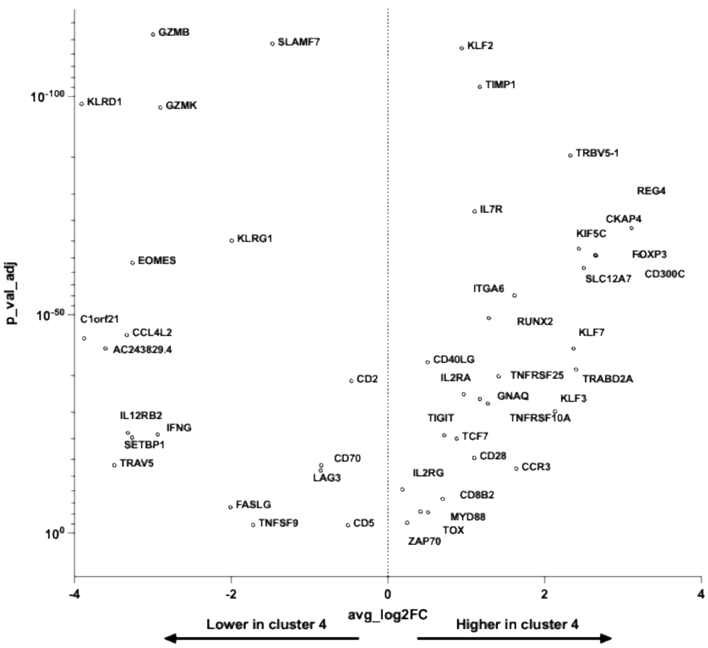


**E**

#### NeoSelect d28 reactivity-seq, selected cluster 5 DEGs

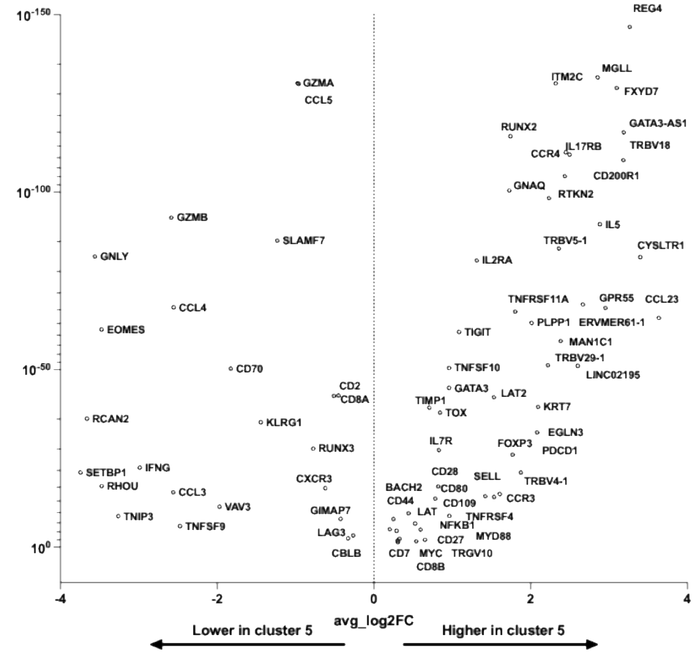


**F**

| NeoTCR | Neoantigen | TRBV | CDR3B | TRAV | CDR3A |
|---|---|---|---|---|---|
| NeoTCR8 #1 | TP53 | TRBV4-1*01 | CASSQGAGNYEQYF | TRAV14/DV4*02 | CAMRERMDSSYKLIF |
| NeoTCR8 #8 | TP53 | TRBV29-1*01 | CSVEGGSGRTGEQYF | TRAV22*01 | CAADYKLSF |
| NeoTCR8 #2 | TP53 | TRBV4-1*01 | CASSQEPRTSGRGEQFF | TRAV38-1*01 | CAFMTGANSKLTF |
| NeoTCR8 #9 | TP53 | TRBV19*01 | CASSIGGSREQFF | TRAV29/DV5*04 | CAEGRADKLIF |
| NeoTCR8 #15 | TP53 | TRBV6-2*01 | CASSYLDYPSSEQFF | TRAV1-1*01 | CAVRGNSGGYQKVTF |
| NeoTCR8 #4 | PSMB9 | TRBV5-1*01 | CASSWGTSESNEQFF | TRAV3*01 | CAVTRGGGNKLVF |
| NeoTCR8 #5 | PSMB9 | TRBV5-1*01 | CASSLQMNYEQYF | TRAV19*01 | CALRSIPQGGSEKLVF |
| NeoTCR8 #10 | PSMB9 | TRBV4-1*01 | CASSLRPDYEQYF | TRAV8-3*01 | CAVEGGSQGNLIF |
| NeoTCR8 #16 | PSMB9 | TRBV5-1*01 | CASSIGTSESNEQFF | TRAV3*01 | CAAPTRGNTGKLIF |
| NeoTCR8 #12 | PSMB9 | TRBV5-1*01 | CASSLGTSEINEQFF | TRAV20*02 | CAVRSTYKYIF |

**G**

| NeoPBL | NeoSelect | TRBV | CDR3B | TRAV | CDR3A |
|---|---|---|---|---|---|
| Clonotype 2 | TP53 | TRBV4-1*01 | CASSQGAGNYEQYF | TRAV14/DV4*02 | CAMRERMDSSYKLIF |
| Clonotype 1 | TP53 | TRBV29-1*01 | CSVEGGSGRTGEQYF | TRAV22*01 | CAADYKLSF |
| Clonotype 6 | TP53 | TRBV27*01 | CASGGWDYGYTF | TRAV12-2*01 | CAVNEAGGYNKLIF |
| Clonotype 7 | TP53 | TRBV6-4*01 | CASSPGTGLLLHF | TRAV13-1*01 | CAAHVGGNTGKLIF* |
| Clonotype 8 | TP53 | TRBV19*01 | CASSAGLAGPLSYEQYF | TRAV13-1*01 | CAASGPNYGGQNFVF |
| Clonotype 14 | PSMB9 | TRBV5-1*01 | CASSWGTSESNEQFF | TRAV3*01 | n/a |
| Clonotype 12 | PSMB9 | TRBV5-1*01 | CASSLQMNYEQYF | TRAV19*01 | CALRSIPQGGSEKLVF |
| Clonotype 13 | PSMB9 | TRBV4-1*01 | CASSLRPDYEQYF | TRAV8-3*01 | CAVEGGSQGNLIF |
| Clonotype 22 | PSMB9 | TRBV5-1*01 | CASSIGTSESNEQFF | TRAV3*01 | CAAPTR_ |
| Clonotype 17 | PSMB9 | TRBV29-1*01 | CSVRTSGSSEQYF | TRAV17*01 | CATVPNSGNTPLVF |
| Clonotype 18 | PSMB9 | TRBV5-1*01 | CASSPGTSDYHNEQFF | TRAV25*01 | CAGRRIGFGNVLHC |
| Clonotype 19 | PSMB9 | TRBV12-4*01 | CASSPGLAVQETQYF | TRAV9-2*02 | CALSPHAGGTSYGKLTF |
| Clonotype 20 | PSMB9 | TRBV5-6*01 | CASSGQGLTEAFF | n/a | n/a |

**Supplemental Figure 6. Reactivity-seq of day 28 NeoPBL from patient 4556.** (A) Table of top upregulated genes in the reactivity cluster (PBL_C9) with $\log_2$ fold changes. (B) Table of notable DEGs in PBL_C9 including TRBV6-4, TRBV4-1, TRBV5-1 and TRBV29-1. (C) UMAPs showing day 28 NeoSelect clonotypes independently designated as reactive, indeterminant or bystander and projected individually onto the 4556 TIL UMAP. (D–E) Selected DEGs associated with C4 (D) and C5 (E), which are enriched for NeoPBL clonotypes. (F) Table showing 10 neoTCR clonotypes originally predicted from 4556 TIL transcriptome data based on high neoTCR8 scores. (G) Table showing 13 NeoPBL clonotypes raised by NeoSelect. Two orange-shaded clonotypes were TP53 reactivities shared between neoTCRs and NeoPBL; four green-shaded clonotypes were shared PSMB9 reactivities.
