## Supplemental Figure 7 for "Rapid enrichment of progenitor exhausted neoantigen-specific CD8 T cells from peripheral blood"

**Supplemental Figure 7. NeoSelect rapidly elicits high-frequency NeoPBL that are revealed to be dominant TIL clonotypes.**

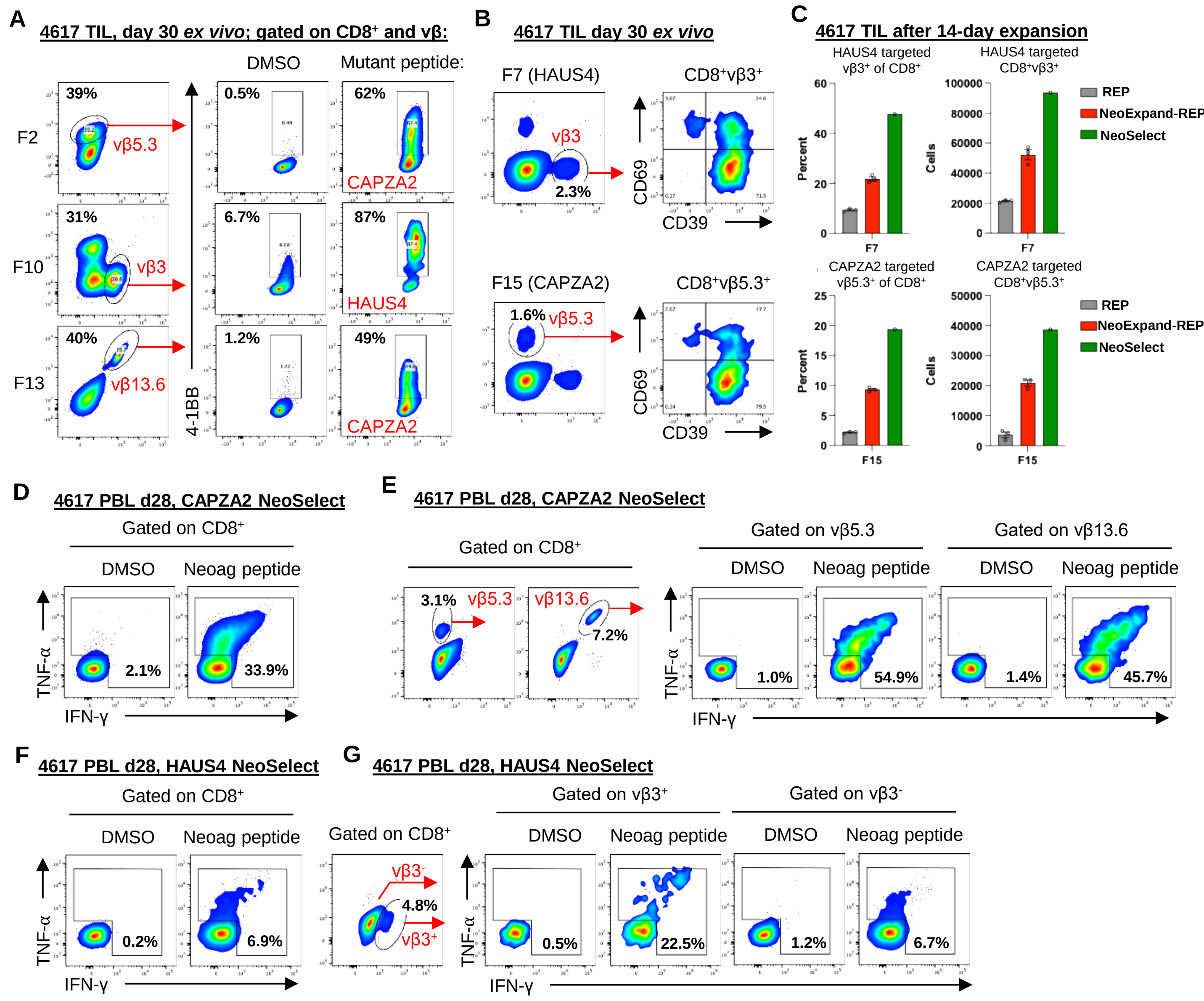


**H** **Patient 4617, sorted CD8+vβ5.3+**

*Ex vivo* cultured TIL

| TCRvβ | CDR3B | Cells | Pct |
|---|---|---|---|
| vβ5.3 | CASSFVRTNTGELFF | 26 | 47.3% |
| vβ5.3 | CASSSPFGQGGFNSPLHF | 14 | 25.5% |
| vβ5.3 | CASMGGHSAPDQETQYF | 3 | 5.5% |
| vβ5.3 | CASRPMDGSNQPQHF | 3 | 5.5% |
| vβ5.3 | CASSFPGGGDNEQFF | 2 | 3.6% |
| vβ5.3 | CASSLGGWFTDTQYF | 2 | 3.6% |
| vβ5.3 | CASSPKVLEGTGMGQNEKLFF | 2 | 3.6% |
| vβ5.3 | CASSLVGRGLVTEAFF | 1 | 1.8% |
| vβ5.3 | CASSPWGGNSPLHF | 1 | 1.8% |
| vβ5.3 | CASSSEQIGQGRNSYEQYF | 1 | 1.8% |

NeoSelect d28 expanded PBL

| TCRvβ | CDR3B | Cells | Pct |
|---|---|---|---|
| vβ5.3 | CASSSEQIGQGRNSYEQYF | 50 | 50.0% |
| vβ5.3 | CASSFVRTNTGELFF | 42 | 42.0% |
| vβ5.3 | CASSLQGDEETQYF | 2 | 2.0% |
| vβ5.3 | CASSPKVLEGTGMGQNEKLFF | 2 | 2.0% |
| vβ5.3 | CASSLRDLRDSPTQETQYF | 1 | 1.0% |
| vβ5.3 | CASSPGLGADTQYF | 1 | 1.0% |
| vβ5.3 | CASSYTSGRYEQYF | 1 | 1.0% |
| vβ5.3 | CSVDPPDSRSGYTF | 1 | 1.0% |

**I** **Patient 4617, sorted CD8+vβ13.6+**

*Ex vivo* cultured TIL

| TCRvβ | CDR3B | Cells | Pct |
|---|---|---|---|
| vβ13.6 | CASSPKVLEGTGMGQNEKLFF | 36 | 65.5% |
| vβ13.6 | CASSFVRTNTGELFF | 12 | 21.8% |
| vβ13.6 | CASMGGHSAPDQETQYF | 2 | 3.6% |
| vβ13.6 | CASRPMDGSNQPQHF | 2 | 3.6% |
| vβ13.6 | CASSSPFGQGGFNSPLHF | 2 | 3.6% |
| vβ13.6 | CVSSSPFGQGGFNSPLHF | 1 | 1.8% |

NeoSelect d28 expanded PBL

| TCRvβ | CDR3B | Cells | Pct |
|---|---|---|---|
| vβ13.6 | CASSPKVLEGTGMGQNEKLFF | 58 | 71.6% |
| vβ13.6 | CASSFVRTNTGELFF | 21 | 25.9% |
| vβ13.6 | CASSSEQIGQGRNSYEQYF | 2 | 2.5% |

**Supplemental Figure 7. NeoSelect rapidly elicits high-frequency NeoPBL that are revealed to be dominant TIL clonotypes.** (A) Flow cytometry plots showing 4-1BB expression in response to DMSO or neoantigen peptide in 4617 TIL fragments, gated on $CD8^+$ T cells and the indicated TCRvβ. (B) Expression of vβ3 and vβ5.3 in fragments 7 and 15, respectively, and CD39/CD69 expression in vβ-gated populations. (C) Frequency of $CD8^+$ T cells expressing vβ3 (fragment 7) or vβ5.3 (fragment 15) after 14 days of REP, NeoExpand-REP, or NeoSelect. (D) Reactivity of $CD8^+$ T cells after 28-day NeoSelect targeting the CAPZA2 neoantigen. (E) Frequency of $vβ5.3^+$ and $vβ13.6^+$ $CD8^+$ T cells and their response to CAPZA2 peptide after 28 days of NeoSelect. (F) Reactivity of $CD8^+$ T cells after 28-day NeoSelect targeting the HAUS4 neoantigen. (G) Frequency of $vβ3^+$ cells and their response to HAUS4 peptide after 28 days of NeoSelect. (H) Tables showing CDR3B sequences obtained from sorting $CD8^+vβ5.3^+$ *ex vivo* cultured TIL (left) or $CD8^+vβ5.3^+$ PBL after 2 cycles of NeoSelect (right). Orange, blue or green shading indicates clonotypes shared between TIL and PBL. (I) Tables as in (H) but after sorting $CD8^+vβ13.6^+$ cells.
